# GD2 and its biosynthetic enzyme GD3 synthase promote tumorigenesis in prostate cancer by regulating cancer stem cell behavior

**DOI:** 10.1101/2023.03.18.533299

**Authors:** Aaqib M. Bhat, Bhopal C. Mohapatra, Haitao Luan, Insha Mushtaq, Sukanya Chakraborty, Siddhartha Kumar, Wangbin Wu, Ben Nolan, Samikshan Dutta, Matthew D. Storck, Micah Schott, Jane L. Meza, Subodh M. Lele, Ming-Fong Lin, Leah M. Cook, Eva Corey, Colm Morrissey, Donald W. Coulter, M. Jordan Rowley, Amarnath Natarajan, Kaustubh Datta, Vimla Band, Hamid Band

**Affiliations:** Eppley Institute for Research in Cancer and Allied Diseases, University of Nebraska Medical Center, Omaha, NE; Department of Genetics, Cell Biology and Anatomy, College of Medicine, University of Nebraska Medical Center, Omaha, NE; Departments of Pathology & Microbiology, College of Medicine, University of Nebraska Medical Center, Omaha, NE; Incyte Corporation, Wilmington, Delaware; Department of Biochemistry and Molecular Biology, University of Nebraska Medical Center, Omaha, NE; Department of Biostatistics, College of Public Health, University of Nebraska Medical Center, Omaha, NE; Department of Urology, University of Washington, Seattle, WA; Department of Pediatrics, University of Nebraska Medical Center, Omaha, NE; Fred & Pamela Buffett Cancer Center, University of Nebraska Medical Center, Omaha, NE

**Keywords:** GD2, GD3 Synthase, prostate cancer, castration-resistant prostate cancer, metastasis, cancer stem cells

## Abstract

While better management of loco-regional prostate cancer (PC) has greatly improved survival, advanced PC remains a major cause of cancer deaths. Identification of novel targetable pathways that contribute to tumor progression in PC could open new therapeutic options. The di-ganglioside GD2 is a target of FDA-approved antibody therapies in neuroblastoma, but the role of GD2 in PC is unexplored. Here, we show that GD2 is expressed in a small subpopulation of PC cells in a subset of patients and a higher proportion of metastatic tumors. Variable levels of cell surface GD2 expression were seen on many PC cell lines, and the expression was highly upregulated by experimental induction of lineage progression or enzalutamide resistance in CRPC cell models. GD2^high^ cell fraction was enriched upon growth of PC cells as tumorspheres and GD2^high^ fraction was enriched in tumorsphere-forming ability. CRISPR-Cas9 knockout (KO) of the rate-limiting GD2 biosynthetic enzyme GD3 Synthase (GD3S) in GD2^high^ CRPC cell models markedly impaired the *in vitro* oncogenic traits and growth as bone-implanted xenograft tumors and reduced the cancer stem cell (CSC) and epithelial-mesenchymal transition (EMT) marker expression. Our results support the potential role of GD3S and its product GD2 in promoting PC tumorigenesis by maintaining cancer stem cells and suggest the potential for GD2 targeting in advanced PC.

## INTRODUCTION

Despite gains from early diagnoses, androgen derivation therapy (ADT) and androgen receptor (AR) signaling inhibitors (ARSIs), prostate cancer (PC) remains a major killer of men and the low survival of patients with distant metastases remains a major clinical challenge. A key transition towards less controllable and incurable metastatic disease is the emergence of castration-resistant PC (CRPC). This has prompted a search for novel CRPC-associated molecular pathways. Molecular targets against which agents with acceptable clinical safety profiles exist are particularly attractive as their transition to clinical translation can be expected to be swifter.

The cell-surface di-ganglioside GD2 is expressed on developing neuroectodermal cells and at lower levels in certain adult tissues like neurons and basal cells of the skin [1]. Biochemically, GD2 is a b-series ganglioside generated from lactosyl-ceramide, which is serially converted to GM3, GD3 and GD2 by the enzymes *ST3GAL3* (GM3 synthase; GM3S), *ST8SIA1* (GD3 synthase; GD3S) and *B4GALNT1* (GM2/GD2 synthase; GD2S), respectively, with GD3S considered rate-limiting [2] (**Fig. 1A**). GD2 is highly expressed in nearly all neuroblastomas, most melanomas and retinoblastomas, and many Ewing sarcomas; it is also overexpressed in some cases of small cell lung cancer, gliomas, osteosarcomas, and soft tissue sarcomas [3].

**Fig. 1:**
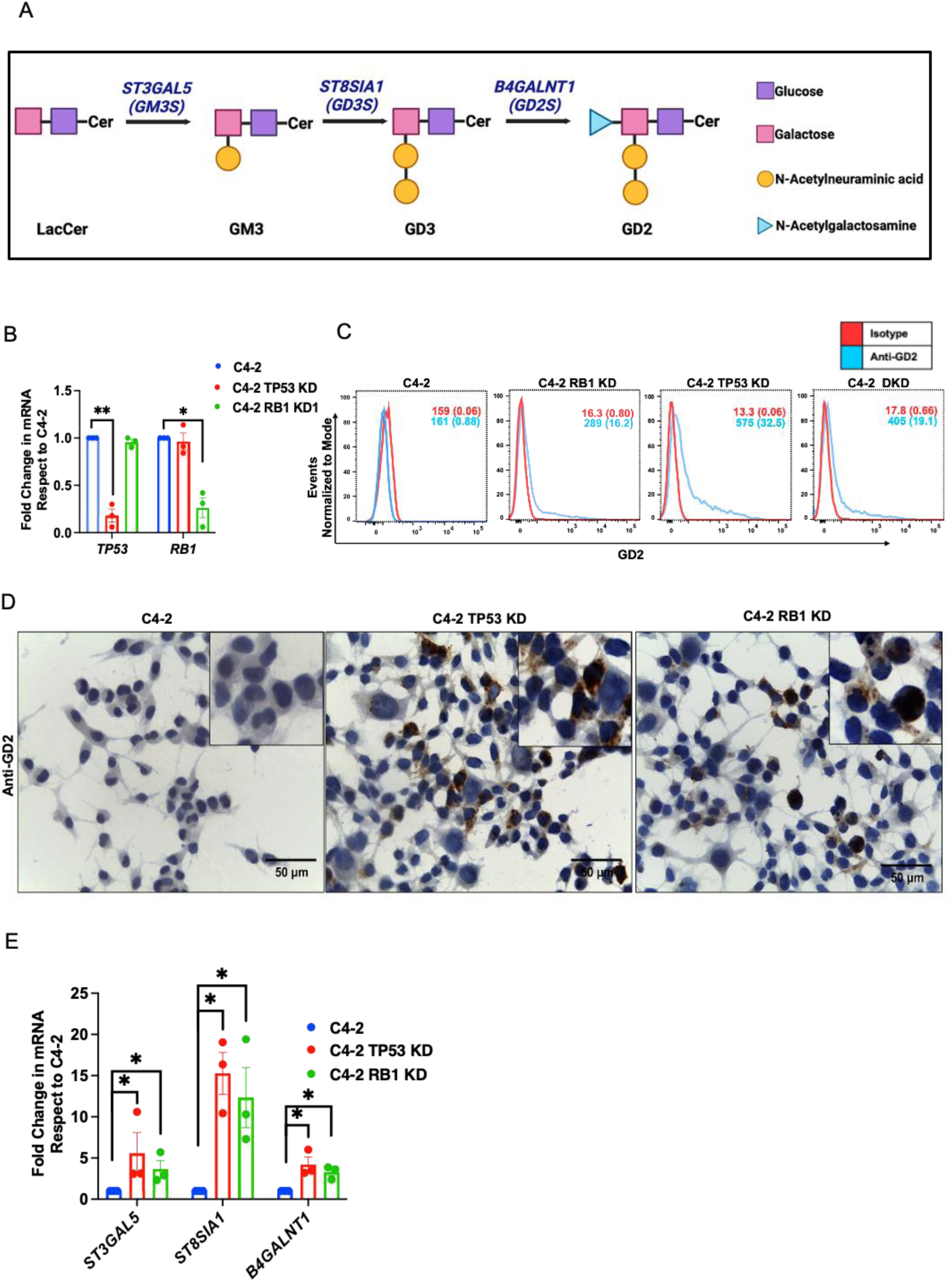
Induction of high GD2 expression in of GD2 in CRPC cell line models with neuroendocrine (NE) differentiation. **(A)** Schematic of key GD2 biosynthetic steps displaying the enzymes (*ST3GAL3, ST8SIA1* and *B4GALNT1*) and intermediate products (GM3 and GD3) [Adapted from [52]]. C4-2 cells with stable knockdown (KD) of TP53 or RB1, previously shown to undergo NE differentiation, were analyzed for GD2. **(B)** qPCR analysis for *TP53* and *RB1* mRNA expression shows TP53-KD or RB1-KD cells relative to C4-2 cells. *n* = 3. **(C)** Cell surface GD2 expression of C4-2, C4-2 RB1-KD, TP53-KD and RB1/TP53 double knockdown (DKD) cells were analyzed by live cell surface staining with anti-GD2 vs isotype control antibodies followed by flow cytometry. X-axis, fluorescence intensity; Y-axis, Events normalized to mode. FACS plots display % cells staining positive and mean fluorescence intensity (MFI) of staining with anti-GD2 (blue) and isotype control (red). **(D)** Immunohistochemistry (IHC) staining confirmation of increased GD2 expression on TP53-KD or RB1-KD C4-2 cells relative to parental cells. **(E)** qPCR analysis shows increased expression of *ST3GAL3, ST8SIA1* and *B4GALNT1 mRNAs* in TP53-KD *and* RB1-KD cells compared to parental C4-2 cell line. *n* = 3. Data are mean ± SEM with two-tailed unpaired *t* test. *, *P* < 0.05; **, *P* < 0.01.

Functionally, GD2 has been implicated in oncogenic signaling leading to increased cell proliferation, migration and invasion, and has been designated by the NCI as the top 12^th^ among 75 tumor antigens [4]. Extensive studies have led to the approval of GD2-targeted antibody Dinutuximab for maintenance therapy of neuroblastoma patients [5, 6], with clinical studies demonstrating the safety and efficacy of this approach [7]. A humanized anti-GD2 monoclonal antibody Naxitamab was recently approved for children and adults with relapsed or refractory high-risk neuroblastoma [8].

Successful GD2 targeting in neuroblastoma has prompted studies of GD2 in other cancers [3]. Significant insights have come from studies of breast cancer. Unlike neuroblastoma, where a majority of tumor cells express GD2, human breast cancer cell lines and patient samples harbor a small GD2^high^ fraction which selectively forms tumorspheres *in vitro* and tumors when implanted *in vivo* and was found to be enriched in triple-negative breast cancer (TNBC), a subtype known to be more aggressive and metastasis-prone [9, 10]. The GD2^+^ breast cancer cells were found to reside in the CD44^hi^/CD24^lo^ cancer stem cell (CSC) fraction, and GD3S knockdown reduced the CSC population and CSC-associated properties and abrogated tumor formation *in vivo* [9, 10]. Further, GD3S knockdown or inhibition of its expression by triptolide inhibited the EMT and migratory/invasive behavior *in vitro* and abrogated metastasis *in vivo* in claudin-low (mesenchymal) breast cancer cell lines [11]. Finally, Dinutuximab inhibited the experimental triple-negative breast tumor growth by targeting CSCs [12]. Recently, O-acetyl-GD2 was also found to mark breast cancer CSCs [13]. GD2 expression on breast cancer CSCs also allowed synergistic targeting by human *γδ* and CD8^+^ T cells together with humanized anti-GD2 antibody hu14.18K322A [14]. Further, GD2-CAR expressing T cells inhibited the orthotopic breast cancer xenograft growth and completely prevented lung metastasis by eliminating CSCs [15]. GD2 expression also correlated with CSC and EMT marker expression in bladder cancer cells [16]. GD3S and GD3 [17] as well as O-acetyl-GD2 [17] were found to drive glioblastoma CSCs and tumorigenicity, with anti-O-acetyl-GD2 antibody sensitization of tumor cells to temozolomide *in vitro* and *in vivo*. Thus, GD2 and its rate-limiting biosynthetic enzyme GD3S have emerged as key factors in promoting CSC behavior and as targets of CSC-directed therapeutic efforts [18, 19]. As CSCs are key to initiation and maintenance of tumors, and responsible for therapeutic resistance and metastasis [20], targeting GD2 on this tumor cell subpopulation represents a novel paradigm about the role of this ganglioside and its enzymatic machinery in non-neuroblastoma tumors.

Progression to CRPC is associated with increased CSC features and EMT [21]. However, few, mostly old, studies have explored the status of GD2 in PC. A study of a small number of prostate hyperplasia, prostate cancer, and normal prostate tissue samples by thin layer chromatography (TLC) found reduced GD2 levels during tumorigenesis [22]. Another chromatography study detected higher GD2 expression in AR-negative PC cell lines (HH870, DU145 and PC-3) than in AR^+^ cell lines (LNCaP clones FGC and FGC-10), and immunofluorescence analyses showed the expression of cell surface GD2 on PC3 cells but not a normal prostate epithelial cells [23, 24]. A recent study found higher expression of a long non-coding RNA (lncRNA), the MIR4435-2HG, in PC cell lines (VCAP, LNCaP, DU145 and PC-3) compared to a normal prostate epithelial cell line, and its knockdown reduced while ectopic overexpression increased the PC3 cell proliferation, migration and invasion *in vitro* and tumorigenesis *in vivo* [25]. MIR4435-2HG was found to bind to GD3S mRNA and to promote its expression, and GD3S knockdown (which by itself had no impact) was found to reduce pro-tumorigenic effects of ectopic MIR4435-2HG overexpression [25]. This study did not examine the ganglioside products of GD3S, and reported high GD3S expression in all 4 PC cell lines examined; the latter is not confirmed by our analysis of the mRNA expression of the same cell lines reported in a comprehensive epigenetics/transcriptomics study [26] and our analyses demonstrate little or no GD2 expression on some of these PC cell lines (See Results). GD2 expression in PC at a histological level and any functional role(s) of GD2 in PC remain unknown. Here, we carry out analyses of GD2 expression in prostate cancer patient tissues and their metastases as well as in cell line models of CRPC progression and use CRISPR-KO of GD3S in human and murine CRPC cell line models to establish that GD2 expression is a feature of a small fraction of tumor cells enriched in tumorsphere growth *in vitro*, and that GD3S is required for tumorigenesis in CRPC.

## MATERIALS AND METHODS

### Prostate cancer (PC) patient tissue microarrays and immunohistochemical analysis

All methods and experimental protocols were carried out in accordance with relevant guidelines and regulations of Institutional Biosafety Committee of the University of Nebraska Medical Center. Tissue microarrays (TMAs) composed of 320 cases of paired normal prostate and prostate cancer Gleason grades (3, 4 and 5) and or of 45 cases of bone and visceral metastatic samples from Rapid Autopsy TMAs were obtained without any identifying patient information from the Prostate Cancer Biorepository Network (PCBN) repository under a materials transfer agreement. 320 prostate specimens used for immunohistochemical analysis were radical prostatectomy samples selected from the surgical pathology files at the Johns Hopkins Department of Pathology with Institutional Review Board approval. 45 cases of metastatic TMAs were obtained from patients who died of metastatic CRPC and signed written informed consent for a rapid autopsy to be performed within hours of death, under the aegis of the prostate cancer Donor Program at the University of Washington. The use of metastatic TMAs for IHC analysis was approved by the Institutional Review Board at the University of Washington.

TMAs were stained for GD2 by immunohistochemistry (IHC) as described previously [27]. Briefly, TMAs were deparaffinized in xylene, rehydrated in descending alcohols and heated in the antigen retrieval solution (vector, #H-3300) in a microwave to unmask the antigens. Endogenous peroxidase activity was blocked by incubation in 3% hydrogen peroxide for 20 min. The sections were rinsed in phosphate buffered saline (PBS) and blocked in Protein-Block buffer (DakoCytomation #0909) for 15 min. The sections were then stained with anti-GD2 antibody (2 μg/ml) overnight at 4°C. After rinsing in TBST, the sections were incubated for 15 min with an anti-mouse secondary antibody conjugated to a dextran-labeled polymer and HRP (DakoCytomation K4007), followed by incubation in DAB solution (DakoCytomation DAB # K4007, Solution 3a, b) for 5 min. The sections were counter-stained with hematoxylin and mounted under cover glasses using a xylene-based mounting medium. GD2 scoring was evaluated by a certified pathologist (SL) and staining scores were assigned as: 0 (negative), 1 (low positive), 2 (moderate positive) and 3 (strong positive). Four cores per case was analyzed for primary tumors and three cores per case for metastasis. Histoscore was calculated by multiplying the staining intensity by % cells staining. Histoscore distributions were assigned as follows: 0-5, negative; 5-25, low; 25-50, moderate; and >50, strong positive. Histoscore 5 was used as a cut off.

### Analysis of *ST8SIA1* mRNA expression in transcriptomic data of CRPC patient-derived xenograft (PDX) tumors and organoid models

*ST8SIA1* mRNA expression was analyzed using publicly available transcriptomic data on CRPC PDX and organoid models published by Tang et al., [26] (GEO accession GSE199190). As per the author classification, the CRPCs were grouped into 4 groups: Group 1, Androgen Receptor dependent (CRPC-AR); Group 2, Wnt dependent (CRPC-WNT); Group 3, neuroendocrine (CRPC-NE); and Group 4, stem cell like (CRPC-SCL) were analyzed. Gene expression is presented as normalized transcripts per million (TPM).

### Cell lines and medium

22Rv1 (ATCC, Cat# CRL-2505) and RM-1 (ATCC, Cat# CRL-3310) cell lines were obtained from ATCC and cultured in RPMI-1640 (Hyclone; #SH30027.02) or DMEM (Gibco; #11965-092), respectively, with 10% fetal bovine serum (Gibco; #10437-028), 10 mM HEPES (Hyclone; #SH30237.01), 1 mM each of sodium pyruvate (Corning; #25-000-CI), nonessential amino acids (Hyclone; #SH30238.01), and L-glutamine (Gibco; #25030-081), 50 μM 2-ME (Gibco; #21985-023), and 1% penicillin/ streptomycin (#15140-122; Gibco). HEK-293T cells (ATCC CRL-3216) were cultured in complete DMEM medium. C4-2 parental, TP53 stable-KD, RB1-KD cell lines, as well as the parental C4-2B and its enzalutamide resistant derivative line (C4-2BER) were grown as described previously [28]. 9464D and 975A2 mouse neuroblastoma cells were kind gifts from Dr. Carrol Thiele, NIH, Bethesda and cultured in DMEM media. Cell lines were maintained for less than 30 days in continuous culture and were regularly tested for mycoplasma.

### Reagents and Antibodies

Primary antibodies used for immunoblotting were as follows: anti-ST8SIA1/GD3S from Novus Biologicals (#NBP1-80750) or ProteinTech (#24918-1-AP); anti-SOX2 from ProteinTech (#11064-1-AP); anti-SOX9 from Abcam (#ab185966); anti-Snail from CST (#3879); anti-HSC70 (#sc-7298); anti-Notch1 (#sc-376403) from Santa Cruz Biotechnology; anti-RB1 (BD Bioscience, #554136); anti-β-actin (#A5441) from Sigma; Anti-Stat3 (#9139S) from CST; anti-Lin28b (#ab191881 from Abcam); anti-Zeb1 (#3396S) from CST; anti-Slug (#ab27568) from Abcam, anti-N-cadherin (#13116S) from CST; anti-CHGA (#MA5-17056) from Invitrogen; anti-NSE (#ab79757) from Abcam and anti-EZH2 (#5246S) from CST. The horseradish peroxidase (HRP)-conjugated Protein A (#101023) and HRP-conjugated rabbit anti-mouse secondary antibody (#31430) and HRP conjugated Goat anti-mouse IgM (#31440) for immunoblotting were from Thermo Fisher. APC conjugated anti-human ganglioside GD2 antibody (#357306; Biolegend) or APC-Mouse-IgG2a,κ isotype control (#400220; Biolegend) was used for IHC.

### Generation of CRISPR/Cas9 knockout and luciferase reporter cell lines

The following CRISPR/Cas9 All-in-One Lentiviral sgRNA constructs were used to generate GD3S knockout (KO) lines. For mouse RM-1, mouse *St8sia1* sgRNA CRISPR/Cas9 All-in-One Lentivector (pLenti-U6-sgRNA-SFFV-Cas9-2A-Puro; #456551140595, Applied Biological Materials) with three target sequences Target 1-17: GGGCCCTACATACGTCCAGA (sgRNA1), Target 2-56: CTCGGAAATTCCCGCGTACC (sgRNA2), Target 3-141: CTACCGGCTGCCCAACGAGA (sgRNA3). For human 22Rv1 and C4-2BER cells, GSGH1195-Edit-R Human All-in-one Lentiviral sgRNA targeting *ST8SIA1* gene, #GSGH1195-247582548 and #GSGH1195-247725854 (designated as sgRNA2 and sgRNA3 respectively) were from Horizon Discovery. Lentiviral supernatants generated in Lenti-x HEK-293T cells (TAKARA, #632180) using the 2^nd^ generation packaging system (plasmids psPAX2, Addgene #12260 and pMD2.G, Addgene #12259) were applied to cells for 72h with polybrene (10 µg/ml, Sigma #H9268) and stable lines selected in 2 μg/ml puromycin. Clones obtained by limiting dilution were analyzed by western blotting for GD3S and FACS for GD2. 3 clones for each sgRNA, maintained separately, were pooled for experiments. The tdTomato-luciferase expressing RM-1 cells were generated using the pMuLE lentiviral system kit (Addgen#1000000060), as described [29]. The mCherry-luciferase expressing 22Rv1 derivatives were generated using a lentiviral plasmid (pCDH-EF-eFFly-T2A-mCherry; Addgene #104833). Cell lines with high reporter expression were obtained by FACS sorting.

### Western Blotting

Western blotting of whole cell extracts prepared in Triton-X-100 lysis buffer and quantified using the BCA assay kit (#23225; Thermo Fisher Scientific) was performed as described previously [27]. 50 μg protein aliquots were run on SDS-7.5% polyacrylamide gels, transferred to polyvinylidene fluoride (PVDF) membrane, and immunoblotted with the indicated antibodies.

### Flow cytometry

Cells seeded in regular medium for 48 h were released with trypsin-EDTA (#15400054; LifeTech (ThermoFisher)) and trypsinization stopped with soybean trypsin inhibitor (#17075029; LifeTech (ThermoFisher). Cells were washed thrice in ice-cold FACS buffer (2% FBS, 25mM HEPES in PBS), and live cells stained with APC anti-human GD2 (#357306; Biolegend) (isotype control, APC Mouse IgG2a, k; #400220, Biolegend) or Anti-human/mouse CD49f antibody (#313620, Biolegend) (isotype control, Pacific Blue Rat IgG2a, k; #400527, Biolegend). For GD3 FACS analysis, anti-GD3 (purified anti-GD3 (MEL-1) antibody, #917701 from Biolegend) with isotype control (purified mouse IgG3, k; #401302) and secondary protein G, Alexa Fluor 488 (#P11065) was used. FACS analyses were performed on a BD LSRFortessa X50, BD LSRII or BD LSRII Green instruments and data analyzed using the FlowJo software.

### Trans-well migration and invasion assay

Cells grown in 0.5% FBS-containing starvation medium for 24 hours were trypsinized and seeded at 2 × 10^4^ on top chambers of 24-well plate trans-wells (Corning, cat# 353097 for migration; or Biocoat Matrigel invasion chambers #354480) in 400 μl of growth factor deprived medium. After 3 hours, medium containing 10% FBS was added to lower chambers and incubated for 16 hours. The non-migrated cells on the upper surface of membranes were removed with cotton swabs, migrated cells on the lower surface were methanol-fixed and stained in 0.5% crystal violet in methanol. Six random 10x fields per insert were photographed, and cells counted using the ImageJ software. Each experiment was run in triplicates and repeated three times.

### Cell proliferation assay

Cell proliferation was assayed as described [29], with 250 cells plated per well in 96-well flat-bottom plates in 100 ml medium and an equal volume of the CellTiter-Glo Luminescent Assay Reagent (#G7571; Promega) added at the indicated time-points. Luminescence was recorded using a GloMax® luminometer (Promega).

### Tumorsphere assay

Tumorsphere cultures were done as described [27] with minor modifications. the tumorsphere medium contained DMEM/F12,Thermo Fisher; #1133032) supplemented with 1% penicillin/streptomycin, 4 μg/ml heparin (Stem cell technologies; #07980), 20 ng/ml Animal-Free Recombinant Human EGF (Peprotech; #AF-100-15), 10 ng/ml Recombinant Human FGF-basic (Peprotech; #100-18B), 1X N-2 supplement (Gibco; #17502-048), 1X B27 supplement (Gibco; #17504-044) and 4% Matrigel (BD Biosciences; #356234]. 2×10^4^ cells were seeded per well in 800 μl volume in ultra-low attachment 24-well plates. Tumorspheres were imaged under a phase contrast microscope after 1 week and quantified using the Image J software. All experiments were done in triplicates and repeated 3 times. We have performed tumorsphere assay using above mentioned tumorsphere medium without Matrigel. Imaging and quantitation methods were same.

### Intra-tibial tumorigenic growth and IVIS Imaging

This study was performed in accordance with relevant guidelines and regulations. All animal experiments were performed in accordance with the ARRIVE guidelines with the approval of the UNMC Institutional Animal Care and Use Committee (IACUC; Protocol Approval # 21-004-06 FC). 4×10^4^ (in 20 μl cold PBS) parental (control) or GD3S-KO (sgRNA1 and sgRNA2) RM-1 cells engineered with lentiviral tdTomato-luciferase or 1×10^5^ (in 40 μl cold PBS) parental (control) or GDS3-KO (sgRNA1) 22Rv1 cells engineered with lentiviral mCherry-enhanced luciferase were injected into tibias of 8-weeks old castrated male C57BL/6 (for RM-1 model) or athymic nude (for 22Rv1 model) mice, respectively, and tumor growth was monitored by bioluminescence imaging. Mice were imaged weekly and followed until the control group reached pre-determined humane end points requiring euthanasia. At necropsy, hind limbs, lungs, spleen, and livers were harvested. For bioluminescence imaging, mice received 200 μl D-luciferin (15 mg/ml; Millipore Sigma #L9504) intraperitoneally 15 min before isoflurane anesthesia and were placed dorso-ventrally in the IVIS™ Imaging System (IVIS 2000). Images were acquired using the IVIS Spectrum CT and analyzed using the Living Image 4.4 software (PerkinElmer). The fold change in total Photon flux (p/s) over time was used to compare tumor growth between the groups.

### Bone morphometry analysis

The harvested tibias were fixed in fresh 4% formalin for a day, transferred to 70% ethanol and scanned using a micro-CT scanner (Skyscan 1172, Bruker) (micro-CT scanning core facility, UNMC). The parameters were 55 kV, 593 181 μA, 0.5 mm aluminum filter, 9 μm resolution, 4 frames averaging, 0.4 rotation step, 180° scanning. The images were reconstructed using the NRecon software (version 1.7.4.6, Bruker microCT), registered and realigned using the DataViewer software (version 1.5.6.2, Bruker microCT) and analyzed using the CTAn software (version 1.18.8.0, Bruker microCT) for the percentage of bone volume (BV/TV), trabecular thickness (Tb.Th), trabecular number (Tb.N) and trabecular separation (Tb.Sp).

### RNA isolation and Real Time qPCR analysis

Total RNA extracted using the TRIzol^®^ Reagent (Ambion, Life Technology) was converted into cDNA with the Invitrogen SuperScript III system (18080-051) and subjected to real-time qPCR using the SYBR Green labeling method (Applied Biosystems; 4309155) on an Applied Bioscience QuantStudio thermocycler. The primer sequences used were: *ST8SIA1:* (forward) 5′-TCCCAGCATAATTCGGCAA-3′ reverse, 5′-ATCTGACAGTGTATAATAAACCCTC-3′); *B4GALNT1* (forward, 5′-TTCACTATCCGCATAAGACAC-3′; reverse, 5′-GTAACCGTTGGGTAGAAGC-3′); *ST3GAL3* (forward, 5′-ACATTGCTTGTGTTTGGAG-3′; reverse, 5′-GGACGACATTCCTTCTGC-3′); *TP53* (Forward, 5’-CCTCACCATCATCACACTGG-3’; reverse, 5’-CACAAACACGCACCTCAAAG-3’); RB1 (Forward, 5’-CTCTCACCTCCCATGTTGCT-3’; reverse, 5’-GGTGTTCGAGGTGAACCATT-3’); *GAPDH (*forward, 5’-CAGAACATCATCCCTGCCTCT GAPDH; reverse, 5’-GCTTGACAAAGTGGTCGTTGAG GAPDH). The fold change in expression was calculated relative to control (parental) cells using the ΔΔCt method and normalized to GAPDH reference gene expression.

### RT^2^ Profiler PCR Arrays

mRNA was isolated from parental and GD3S-KO RM-1 cells using QuantiTech Reverse Transcription Kit (#205310). The expression of specific genes was quantitated by qPCR using the RT^2^ Profiler PCR Array (PAMM-176ZA-6 RT^2^ Profiler^TM^ PCR Array for mouse cancer stem cells; catalog #330231) with QuantiTech SYBR Green PCR Kit (#204141) on an Applied Bioscience QuantStudio thermocycler. The analysis was performed using the GeneGlobe online application (geneglobe.qiagen.com) and the fold regulation of the genes was represented against the control in the heatmap and volcano plot.

### Isolation of Gangliosides, Thin Layer Chromatography, and Immunostaining

The isolation of gangliosides, thin layer chromatography and immunostaining were carried out using previously described protocol with minor modifications [30]. 8 × 10^7^ cells were trypsinized and resuspended in 2 ml of Chloroform: Methanol (at the ratio 2:1, v/v), and sonicated in an ultrasonic bath (Emerson Branson 1800) for 15 min at room temperature. The cells were centrifuged @ 2000 rpm for 5 min and supernatants were transferred to clean glass tube with PTFE-lined screw cap. Re-extraction of lipids was performed by adding 2 ml of Chloroform: Methanol (1:1, v/v) and sonicated again for 15 min. The cells were centrifuged again @ 2000 rpm for 5 min and supernatant transferred to clean glass tube. Final extraction of lipids was done by adding 2 ml of Chloroform: Isopropanol: Water (7:11:2 ratio) and sonication for 15 min. The cells were centrifuged again @ 2000 rpm for 5 min and all supernatants were pooled together into a clean glass tube. C18 solid phase extraction cartridge (Waters, SepPak Plus long C18, #WAT023635) was prepped by first washing with 3 mL of methanol, then with 3 ml of Chloroform: Methanol: Water (2:43:55, v/v/v). The pooled supernatant was loaded onto the washed C18 cartridge using the glass syringe and the flow-through collected and reloaded onto the column to optimize adsorption. This was followed by washing the column with 5 ml water to desalt. The gangliosides were eluted with 3 ml of methanol, collecting the eluate in a fresh glass tube and the solvent was removed using Rota-Vapor R-210. The sample was dissolved in 200ul methanol and were run for Thin Layer Chromatography (TLC).

The coated Silica Gel-60 TLC plates (Sigma #HX30882453) were pre-run in chloroform to eliminate neutral lipid and other contaminants that may interfere with the mobility of gangliosides. The samples were loaded on to the dried TLC plates using Hamilton Syringe 1cm above the bottom and 1 cm apart from each other. The loaded samples were air dried and then run in the solvent Chloroform: Methanol: 0.2% CaCl_2_ (5:4:1, v/v/v) [Have tested 55:45:10, 45:55:10, 60:35:10, 70:30:10, 80:20:10]. The TLC plate after the run was allowed to air dry and visualized using Phosphomolybdic acid [12-Molybdophosphoric acid in Ethanol (10% v/v)]. 1 μl (1mg/ mL conc) of Ganglioside, GD2 standard (Cat#25487, Cayman Chemical) was loaded as a positive control. For immunostaining, the TLC plate after the run was dip in 0.2% polyisobutyl-methacrylate Hexane: Chloroform (9:1, v/v) for 1 min with sample side up and dried. The TLC plate was blocked using 1% BSA in PBS for 30 mins at room temperature followed by 3 washes with PBS (3 min each). The plate was then immuno-stained overnight with 0.5 mg/ml anti-GD2 antibody at 4°C. The next day the plate was washed 3 times with PBS (3 min each) and stained with HRP conjugated secondary antibody at 1:500 dilution for 1 hr. at room temperature. The plate was washed thrice with PBS and developed using the ECL substrate.

### Statistical analysis

GraphPad Prism software (version 9) was used to perform the statistical analyses. The *in vitro* experimental data analysis used a two-tailed student’s t test. A two-way ANOVA (mixed model) test was used to analyze the fold change in photon flux in xenograft studies. P values equal to or <0.05 were considered significant.

### Data availability

The data generated in this study are available upon request from the corresponding authors.

## RESULTS

### Increased GD2 expression upon RB1 or TP53 knockdown in a CRPC cell line model

Given the GD2 overexpression in neuroectodermal lineage tumors [3], and prior reports that CRPC-NE differentiation is associated with the acquisition of CSC features [31], we surmised that GD2 overexpression might be a feature of CRPC upon experimental perturbations shown to promote neuroendocrine differentiation. RB1 or TP53 depletion in androgen receptor-driven CRPC (CRPC-AR) cell lines [32] or mouse models [33] combined with PTEN loss is known to promote NE trans-differentiation. Hence, we compared the CRPC-AR cell line LNCaP C4-2 (referred to as C4-2), which has a mutationally-inactivated PTEN [34], with its stable TP53, RB1 or TP53/RB1 knockdown (KD) derivatives as previous studies in this model showed that TP53/RB1-KD cells had acquired NE differentiation [28]. qPCR analysis verified the expected TP53 or RB1-KD (**Fig. 1B**). Western blotting confirmed the knockdown of RB1 and TP53 in individual KD and DKD cells (Supplementary Fig. S1A). We validated a commercially available anti-GD2 antibody (clone 14.G2; Biolegend) by demonstrating its reactivity by FACS and immunohistochemistry (IHC) against a known GD2^+^ neuroblastoma cell line SK-N-BE-2 [35] and the lack of its reactivity with a GD2^-^ rhabdomyosarcoma cell line A204 [36] (**Supplementary Fig. S1B,C**). Using this antibody, FACS staining revealed low cell surface GD2 expression on parental C4-2 cells, but the levels of GD2 and the size of the GD2^+^ subpopulation markedly increased in RB1-KD, TP53-KD or double-KD (DKD) C4-2 derivatives (**Fig. 1C**). Notably, the increase in GD2 levels on DKD was intermediate between that with individual KD of RB1 or TP53, the latter being higher than with RB1-KD (the MFI and % GD2^+^ cells are indicated inside FACS plots). IHC analyses confirmed the upregulation of GD2 in RB1-KD and TP53-KD cells compared to the parental C4-2 cells (**Fig. 1D**). Increased expression of GD2 upon TP53 or RB1-KD of C4-2 cells was associated with increased levels of the key GD2 biosynthesis enzymes GM3 Synthase (GM3S), GD3 synthase (GD3S) and GD2 synthase (GD2S) (**Fig. 1 E**).

### Increased GD2 expression in a subset of PC cell lines

Using the C4-2 and its RB1 or TP53-KD derivatives as references, we carried out FACS analysis of cell surface GD2 expression on a panel of 18 established human and murine PC cell lines, which included variants of commonly used PC cell lines and two immortal prostate epithelial cell lines (PHPV18 and PSV40). Among these, a majority displayed little or no anti-GD2 staining above that with the control antibody (**Table 1, Supplementary Fig. S2A-C**). In contrast, low but clearly detectable levels of GD2 expression on a small subpopulation of cells were observed on 3 PC cell lines (ND1, C4-2 and C4-2B; ranging from 0 to 2%) and two immortal prostate epithelial cell lines (PHPV18 and PSV40; 2 to 3% and 4 to 6 %, respectively) (**Table 1, Supplementary Fig. S2A-C**). Importantly, a relatively large subpopulation of GD2^+^ cells displaying high mean fluorescence intensities (MFI’s) was observed in human CRPC cell lines 22Rv1 (% +, 20.1; MFI 870), which overexpresses wildtype AR and ARV7 splice variant [37], and C4-2B cells rendered enzalutamide-resistant [28] (% +, 50.6; MFI 2528), and on murine CRPC cell lines RM-1 (driven by Ras and Myc) [38] (% +, 54.6; MFI 1088) and TRAMP-C1 (transgenic adenocarcinoma mouse prostate [TRAMP]-derived) [39] (% +, 28.6; MFI 908) (**Table 1, Supplementary Fig. S2A-C**).

**Table-1:** Prostate cancer cell lines show % of GD2 on live cell FACS analysis using anti-GD2 staining.

| **Cell Lines** | **% GD2** |
| --- | --- |
| **PC3** | **0** |
| **PC3M** | **0** |
| **PC3M LN4** | **0** |
| **LNCaP (Low Passage)** | **0** |
| **MDA PCa2b (Low Passage)** | **0** |
| **NE1.3** | **0** |
| **DU145** | **0** |
| **ND1** | **~0 - 2** |
| **LNCaP C4-2** | **~0 - 2** |
| **LNCaP C4-2B** | **~0 - 2** |
| **PHPV18** | **~2 - 3** |
| **PSV40** | **~4 - 6** |
| **22Rv1** | **~15 - 30** |
| **LNCaP C4-2 RB1 KD** | **~10 - 20** |
| **LNCaP C4-2 p53 KD** | **~10 - 35** |
| **TRAMP-C1** | **~25 - 60** |
| **LNCaP C4-2BER (Enzalutamide Resistant)** | **~25 - 60** |
| **RM-1** | **~45 - 70** |

The GD3S enzyme (encoded by the *ST8SIA1* gene) is rate-limiting for GD2 synthesis [2]. Consistent with our staining data, the CCLE-DepMap mRNA expression data available for ten PC cell lines showed the highest value for the GD2^high^ 22Rv1 and the second lowest value for the GD2^-^ PC3 cell line (**Supplementary Fig. S3, upper panel)**. Analysis of transcriptomics data of organoids and PDX tumors recently used to classify CRPC into 4 subtypes, CRPC-AR (AR-driven), CRPC-NE (neuroendocrine), CRPC-WNT (WNT pathway-driven) and CRPC-SCL (stem cell like) [26], revealed high *ST8SIA1* mRNA expression in GD2^high^ 22Rv1 and nearly undetectable levels in GD2^-^ LNCaP cell line (**Supplementary Fig. S3, upper panel**). Using the levels in these cell lines as reference, >20% of CRPC organoids and PDX tumors showed higher *ST8SIA1* expression (those above the red cutoff line corresponding to TPM values for LNCaP), and this was seen across the CRPC subtypes (**Supplementary Fig. S3**). Interestingly, the *ST8SIA1* TPM values in a number of organoid models were substantially higher than those for 22RV1 (e.g., CRPC-AR samples MSK-PCa19, MSK-PCa22; CRPC-NE samples MSK-PCa10, MSK-PCa24; and CRPC-SCL sample MSK-PCa8). Thus, our results show that a subset of established PC cell lines, including examples of CRPC, express low to high GD2 levels on a variable fraction of cells, reminiscent of studies with breast cancer cell models [9, 10], and the *ST8S1A1* expression in organoid and PDX models supported the potential of high GD2 expression in patient-derived PCs.

### GD2 expression is seen on a small subset of tumor cells in primary PC patient tissues and a higher proportion of metastatic lesions

To assess if GD2 expression on a subpopulation of cells in PC cell lines could be extended to human PC, we used tissue microarrays (TMAs) from two PC patient cohorts (provided by the Prostate Cancer Biorepository Network; PCBN). IHC staining of a 320-sample TMA of paired normal prostate tissues and PC samples (Gleason scores of 3-5) demonstrated a statistically significant overexpression of GD2 on primary PCs compared to paired normal prostate tissue (**Fig. 2A-D**); unlike uniform GD2^+^ neuroblastomas, but similar to triple-negative breast cancers, only a small subset of tumor cells in PC samples was GD2^+^. Overall, 19% of primary PCs scored GD2^+^ based on a histoscore cutoff of <5 (**Fig. 2E)**. Notably, IHC analysis of a metastatic PC TMA (the 45-Case Bone and Visceral Metastasis from Rapid Autopsy TMA from PCBN) showed GD2^+^ staining in about twice the percentage of metastatic lesions compared to primary tumors (**Fig. 2E**). The histoscore distribution analysis showed that metastatic tumors exhibited higher GD2 expression (**Fig. 2F-G**). These results indicate that GD2-expression is a feature of a subpopulation of tumor cells in a subset of PC patients, especially in metastases.

**Fig. 2:**
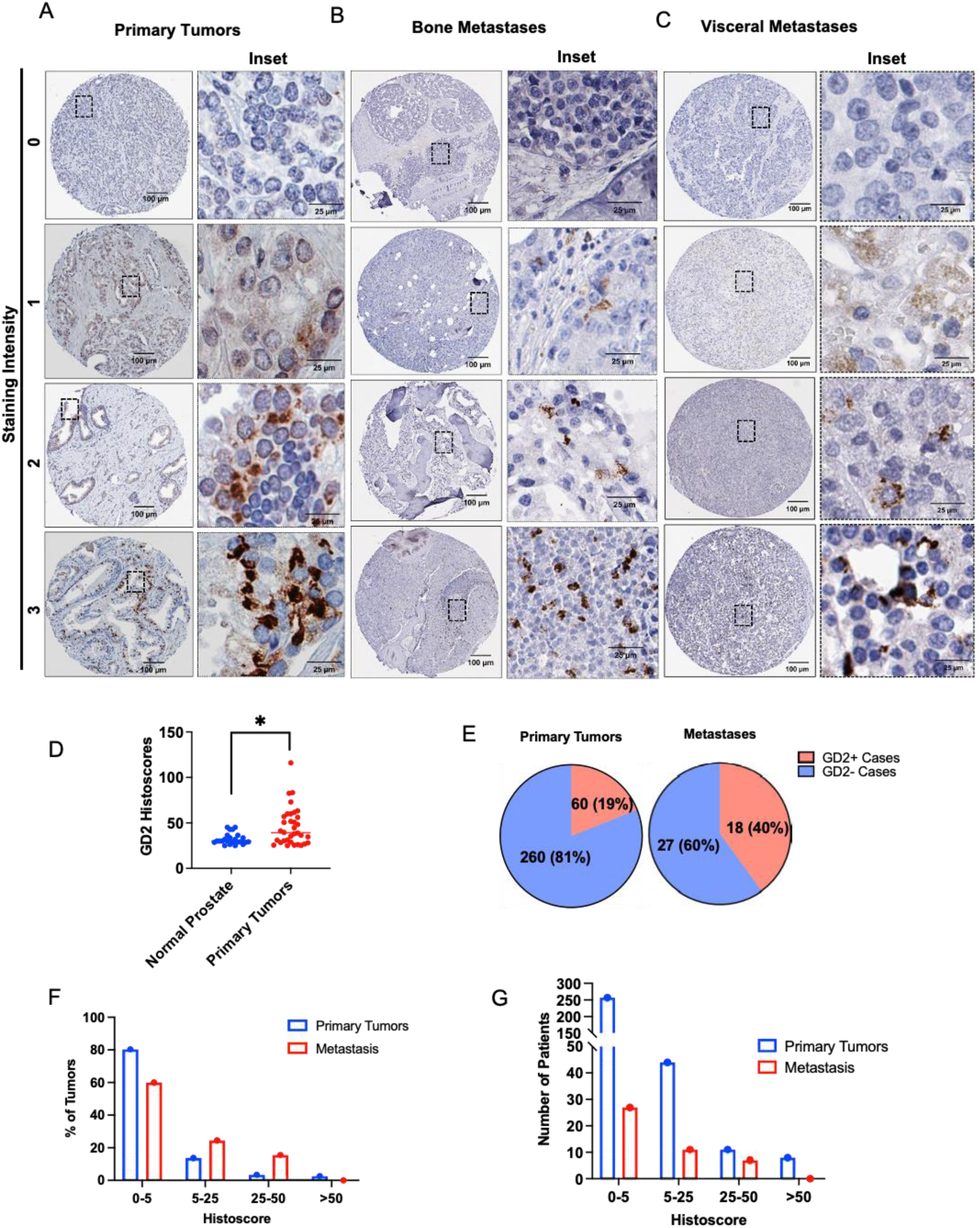
GD2 expression on a subset of tumor cells in primary prostate cancer tissue with higher expression in metastatic lesions. Tissue microarrays (TMAs) composed of paired normal prostate and prostate cancer Gleason grades (3, 4 and 5) (n = 320) or 45 cases of bone and visceral metastatic samples (both obtained from the Prostate Cancer Biorepository Network) were analyzed by IHC staining for GD2. Histoscores were calculated by multiplying the staining intensity (0, negative; 1, low; 2, moderate positive; 3, strong positive) by the % tumor cells staining positive. (**A, B**) Representative examples of negative, low, moderate, and high GD2 staining in **(A)** primary, **(B)** bone metastatic and (**C)** visceral metastatic tumor samples. (**D)** Scattered plot shows histoscore distribution of primary tumors and normal prostate. GD2 expression is significantly higher in primary tumors than normal prostate. Data represents 34 cases of GD2 expression in tumors vs 24 in normal tissue, considering histoscore >25 as positive. Mann-Whitney unpaired t-test used as statistical analysis; *, *P* < 0.05. **(E)** Pie charts displaying the % of cases that were GD2^+^ or GD2^-^ in primary and metastatic tumors samples. Histoscores of 0-5 were considered negative while scores higher than 5 were considered positive. (**F, G**) Frequency distribution plots histoscores plotted against **(F)** percentage of patients and **(G)** number of patients. The indicated histoscore grouping are: 0-5 (negative); 5-25 (low), 25-50 (moderate); and >50 (strong).

### GD2 overexpression defines a PC cell population with higher tumorigenic potential *in vitro*

To assess the potential functional role(s) of GD2 overexpression in PC, we first compared the parental C4-2 cells (GD2^low^) with their GD2^high^ RB1-KD, TP53-KD or DKD cell lines. Cell Titer-Glo assays showed that RB1-KD, TP53-KD or DKD derivatives exhibited higher (fold increase of ∼6, ∼20, ∼31 and ∼23 for parental, RB1-KD, TP53-KD and DKD, respectively; p values <0.01 to<0.001 at the last time point) and faster cell proliferation, with TP53-KD cells, which also exhibited higher GD2^+^ subpopulation, exhibiting the highest rate and level of proliferation (**Fig. 3A, left panel**). Similarly, the GD2^high^ enzalutamide resistant C4-2BER showed increased proliferation compared to GD2^low^ control C4-2B parental cells (fold increase of ∼60, vs ∼11; p value < 0.001 at various time points) (**Fig. 3A, right panel**). Transwell migration analyses showed a higher migratory behavior of GD2^high^ derivatives (∼ 2-fold for RB1-KD, and ∼3.5-fold for TP53-KD and DKD respectively compared to C4-2; ∼4-fold for C4-2BER vs. C4-2B) compared to GD2^low^ parental cell lines (**Fig. 3B**). Importantly, the GD2^high^ derivatives of C4-2 exhibited significantly elevated 3D tumor-sphere growth, an *in vitro* correlate of CSC activity [40], with RB1-KD yielding ∼4.5-fold and TP53-KD or DKD cell lines yielding ∼9-fold more tumorspheres (p-values <0.001) compared to their GD2^low^ parental C4-2 cells (**Fig. 3C, upper panel**). Similarly, GD2^high^ C4-2BER cells exhibited a ∼4.5-fold increase in tumorspheres compared to parental C4-2B cells (p-value <0.001) (**Fig. 3C, middle panel**). Additional tumorsphere assays on ultra-low attachment conditions in the absence of added 4% Matrigel, which we used in above assays to obtain more uniform tumorspheres, confirmed the above findings and excluded the possibility that elevated tumorsphere growth of GD2^high^ C4-2 derivatives was due to adhesion abilities provided by Matrigel (**Fig. 3C, lower panel**).

**Fig. 3:**
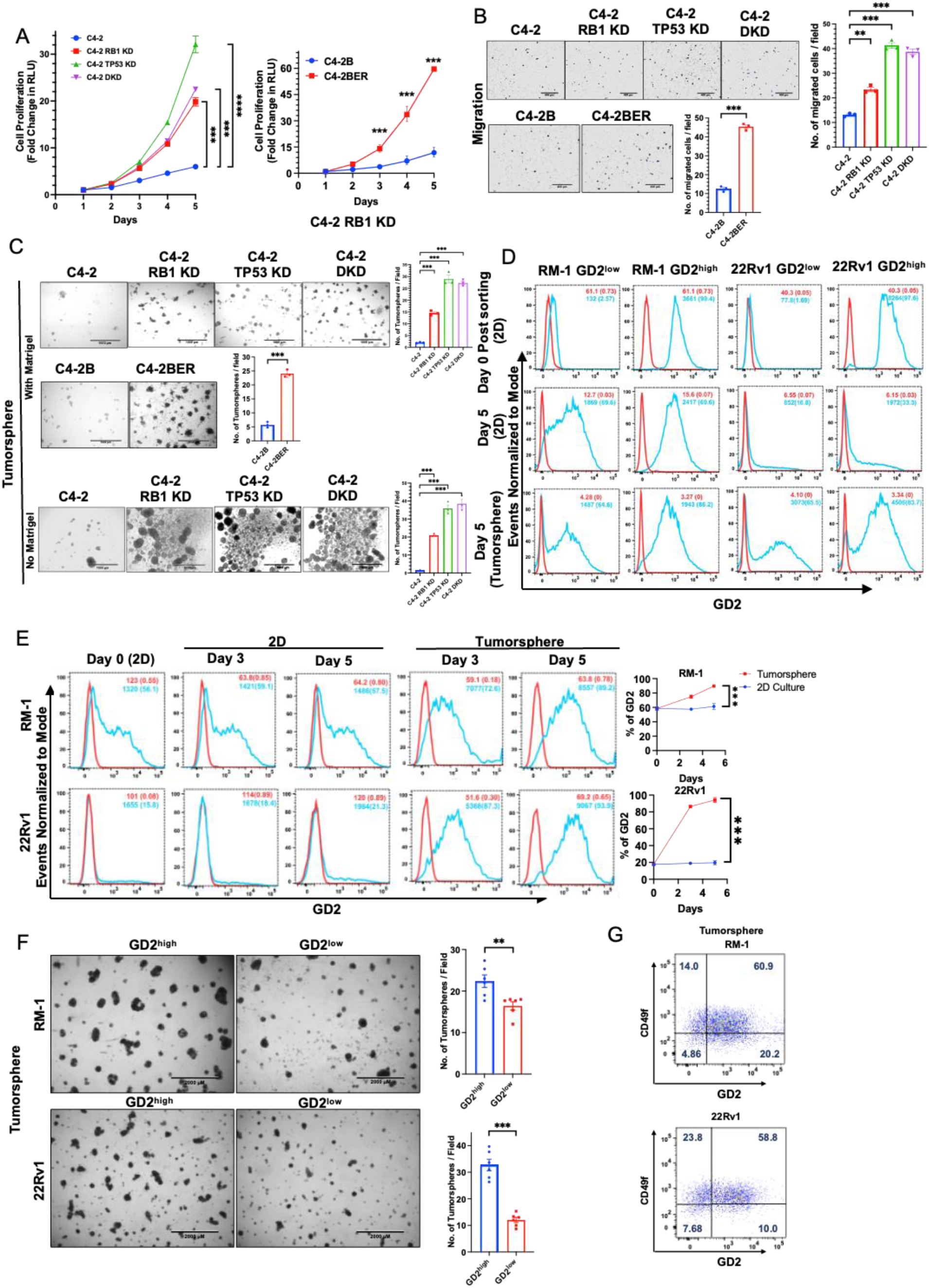
GD2 overexpression promotes the *in vitro* oncogenic traits: (**A**) Analysis of cell proliferation. The parental C4-2 and its TP53-KD or RB1-KD or TP53/RB-KD (DKD) cells and C4-2B and its enzalutamide resistant variant C4-2BER were plated in 96-well plates and cell proliferation was assessed on the indicated days using the CellTiter-Glo luminescent cell viability assay. The Y-axis represents the fold increase in relative luminescence units (RLU) relative to Day 1. *n* = 3 with six replicates each (**B**) Analysis of cell migration in indicated cell lines using 20,000 cells as input. Left, representative images. Right, quantification. *n* = 3 with three replicates each. (**C**) Analysis of tumorsphere forming ability. The indicated cell lines were plated in 24-well ultra-low attachment plates in presence of 4% Matrigel (upper and middle panels) and in absence of Matrigel.(lower panel) and images were obtained on Day 7.Left, representative images; scale bar, 1000 µm. Right, quantified tumorsphere numbers per 4X microscopic field. *n* = 3 with three replicates each and 6 images per well (**D**) GD2^high^ and GD2^low^ population were FACS sorted in RM-1 and 22Rv1 cells and grown as 2D and tumorspheres. After Day 5, the cells were reanalyzed for GD2 fractions in both GD2^high^ and GD2^low^ post sorted cells. In both tumorspheres and 2D culture the GD2^low^ cells re-expressed GD2. (**E**) Enrichment of GD2^high^ cells in tumorspheres of CRPC cell lines. The indicated cell lines were cultured in 2D or were seeded in tumorsphere cultures. Cells harvested at the indicated times were analyzed for cell surface GD2 expression (vs. isotype control) at the indicated times. % GD2^+^ cells and MFI are indicated. Left, representative FACS images; Right, quantitation of data. two-way ANOVA test. (**F**) GD2^high^ fraction of CRPC cell lines is enriched for tumorsphere-forming ability. Constitutively GD2-expressing RM-1 and 22Rv1 CRPC cell lines were FACS sorted for GD2 and GD2 populations, grown on 2D overnight and plated at 20,000 cells per well of 24-well ultra-low attachment plates in tumorsphere media containing 4% Matrigel. Tumorspheres were imaged on Day 5. Left, representative images; scale bar, 2000 µm. Right, Quantification of tumorspheres (>250 µm) per 2X magnification field. *n* = 3 with three replicates each and 6 images per well. (**G**) FACS analysis of tumorsphere-grown RM-1 and 22Rv1 cell lines after anti-GD2 and anti-CD49f double-staining shows a majority of GD2^+^ cells to be CD49f^hi^. % GD2 and/or CD49f +/- populations are shown in the respective quadrants. Data represented in all experiments are mean ± SEM, unpaired t-test; ^ns^, not significant; *, *P* < 0.05; **, *P* < 0.01; ***, *P* < 0.001.

Previous studies have shown that GD2-expressing cell fraction is enriched in tumorigenic behaviors [9]. We therefore used FACS-sorting to purify the GD2^high^ and GD2^low^ populations of 22Rv1 and RM-1 cell lines as these cell lines harbor both GD2^high^ and GD2^low^cell subpopulations (**Table 1**). Consistent with previous studies demonstrating that GD2 expression on cell lines is dynamic [41], the FACS-sorted populations drifted towards a mix of GD2^high^ and GD2^low^ populations upon continued 2D or 3D culture (**Fig. 3D**). Notably, FACS analysis revealed a significant enrichment for the GD2^high^ population during tumorsphere growth vs 2-D culture in both RM-1 (72% and 89% at days 3 and 5 in 3D, and 56% and 59% at 3 and 5 days in 2D, vs. 56% on Day 0; p<0.001) and 22Rv1 (87% and 94% at days 3 and 5 in 3D, and 18% and 21% at 3 and 5 days in 2D, vs. 16% on Day 0; p<0.001) cell lines (**Fig. 3E**). A similar, time-dependent, increase in the GD2^high^ population was observed upon tumorsphere vs. 2D growth of C4-2 (GD2^low^) and its RB1-KD (GD2^high^) derivative (**Supplementary Fig. S4**). Despite the dynamic nature of GD2 expression, the GD2^high^ fractions of both 22Rv1 and RM-1 cell lines yielded significantly more tumor-spheres (∼22/field vs. 16/field for GD2^high^ and GD2^low^ RM-1, and 32/field vs. 11/field for GD2^high^ and GD2^low^ 22Rv1; p<0.01 and <0.001, respectively) (**Fig. 3F**) indicating a higher tumorsphere-forming ability of GD2^high^ subpopulation in PC cell lines. Dual FACS staining of RM-1 and 22Rv1 cell lines grown as tumorspheres for CD49f and GD2 demonstrated that a majority (60.9% for RM-1 and 58.8% for 22Rv1) of GD2^high^ cells were within the CD49f^high^ fraction (**Fig. 3G**). These results suggested that GD2^high^ fraction defines a cell population with higher oncogenic potential and potentially the CSC-like behavior.

### GD3 synthase, the rate-limiting enzyme in GD2 biosynthesis, is required for *in vitro* oncogenic attributes of GD2^high^ CRPC cell lines

In view of our results above, we hypothesized that the GD2^+^ cell subpopulation is the major driver of the oncogenic attributes of PCs. To test this hypothesis, we targeted the rate-limiting GD2 biosynthesis enzyme GD3S [2] for CRISPR-Cas9 knockout (KO), using a single-vector Cas9/sgRNA expression system, in GD2-overexpressing mouse (RM-1) and human (22Rv1) CRPC cell lines and obtained clonal lines by serial limiting dilution cloning. Western blotting confirmed the loss of GD3S expression in GD3S sgRNA-targeted derivatives (**Fig. 4A, B**). These GD3S-KO clones indeed lacked the cell surface GD2 expression by FACS analysis (**Fig. 4C, D**). We also analyzed the parental vs. GD3S-KO RM-1 and 22Rv1 cell lines together with GD2^+^ 9464D and GD2^-^ 975A2 mouse neuroblastoma cell lines [41] by thin layer chromatography (TLC) and TLC/immunoblotting, in which lipid extracts of cells were resolved by TLC next to the purified GD2 standard, followed by either chemical staining to visualize the separated sphingolipid species or immunoblotted with the 14G2a antibody, respectively. Based on pilot studies using various ratios of the chloroform/methanol/0.2% CaCl_2_ solvent system used in prior studies [42-44], we resolved the sphingolipid extracts of cell lines using the 50:40:10 ratio of chloroform/methanol/0.2% CaCl_2_ as these were optimal to resolve GD2 and GD3 species from other lipids. Phosphomolybdic acid (PMA) staining of the resolved species on the TLC plate showed lipid species comigrating with standard GD2 in RM-1, 22Rv1 and the positive control 9464D cells, while these species were absent in the GD3S-KO cell lines and the GD2 negative 975A2 cells (**Supplementary Fig. S5A**). Notably, 14G2a immunoblotting of the TLC-resolved sphingolipid species showed strong reactivity with the GD2 standard, as expected; additionally, the 14G2a antibody detected the lipid species comigrating with standard GD2 in parental RM-1 or 22Rv1 and in the positive control cell line 9464D (corresponding to the comigrating species detected by chemical staining) but not in GD3S-KO RM-1 or 22Rv1 cell extracts or in the negative control cell line 975A2 (**Supplementary Fig. S5B**). These results further confirm the reactivity of 14G2a with GD2 and the absence of its expression upon GD3S-KO. As GD3 is the immediate product of GD3S activity and a precursor of GD2, we also assessed the levels of GD3 in panels of PC cell lines (**Supplementary Fig. S6**). In addition, parental vs. GD3S-KO CRPC cell lines RM-1 and 22Rv1 were assessed for expression of GD3 using FACS. Relatively low to modest levels of GD3 were seen on parental cells and this expression was eliminated in KO cells (**Supplementary Fig. S7**). Thus, GD3S-KO effectively eliminated the expression of GD2 (and GD3) on CRPC cell lines.

**Fig. 4:**
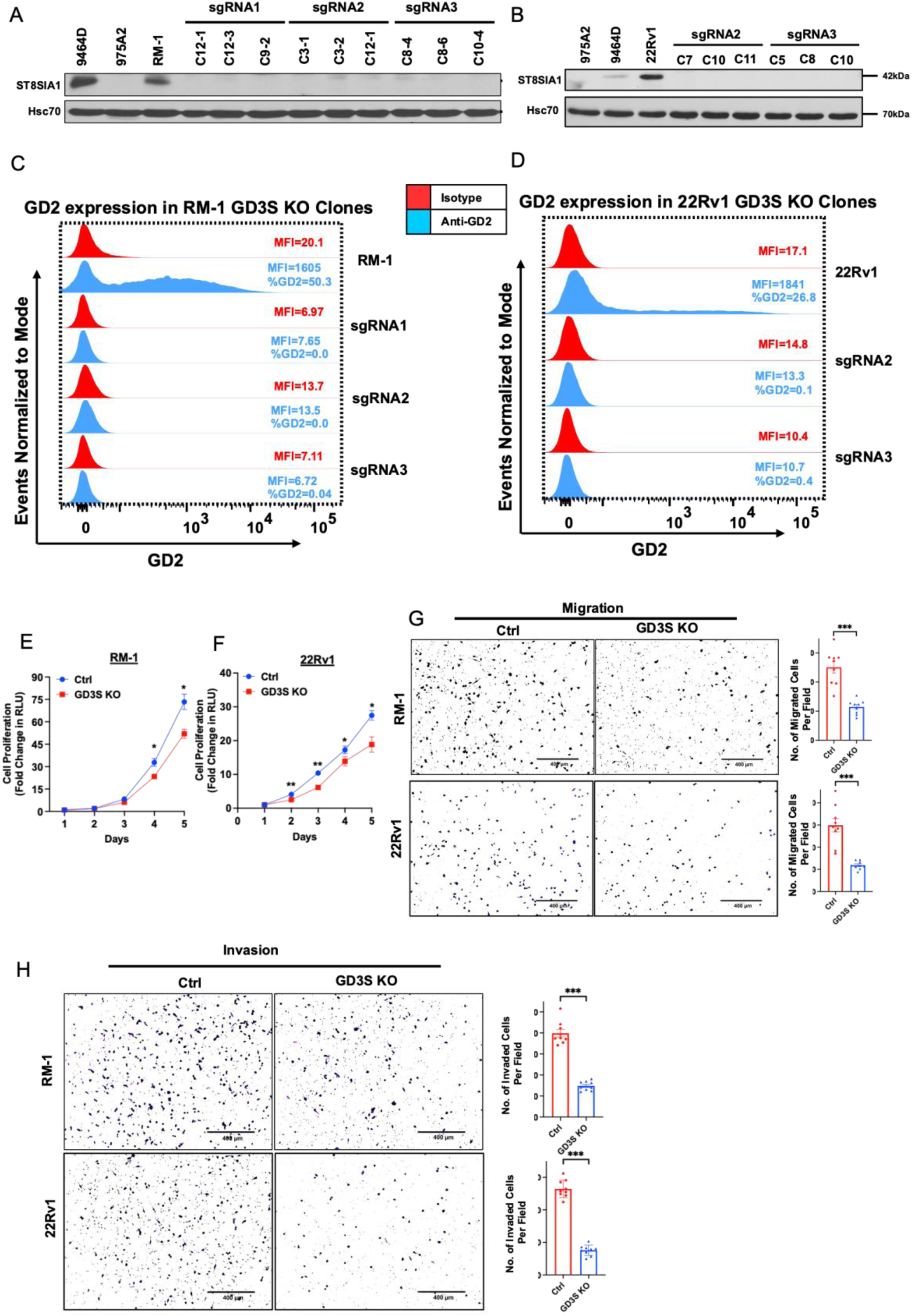
*GD3S* knockout in CRPC cell lines impairs the in vitro tumorigenic and pro-metastatic traits. (**A-D**) Generation of GD3S knockout of RM-1 and 22Rv1 CRPC cell lines. Cells were transduced with lentiviral All-in-One CRISPR/Cas9 constructs and stable clones analyzed by Western blotting with anti-GD3S antibody; Hsc70, loading control. Known GD2^+^ (9464D) and GD2^-^ (975A2) mouse neuroblastoma cell lines were used as controls. (**C & D**) Complete loss of cell surface GD2 expression in GD3S-KO RM-1 and 22Rv1 clones. The parental cell lines and their GD3S-KO clones were live-cell stained with anti-GD2 (blue) or isotype control (red) antibody followed by FACS analysis. FACS plots indicate mean fluorescence intensity (MFI) on X-axis vs. Events Normalized to Mode on Y-axis. % positive cells and MFIs are indicated in FACS plots. (**E & F**) Reduced CRPC cell proliferation upon GD3S-KO. Three clones of parental and GD3S-KO cell lines maintained separately were mixed plated at 1,000 cells per well in 96-well plates and live cells quantified at the indicated times using the CellTiter-Glo cell viability assay. The Y-axis represents the fold increase in relative luminescence units (RLU) relative to Day 1. *n* = 3 with six replicates each. (**G & H**) Reduced CRPC cell migration (**G**) and invasion (**H**) upon GD3S-KO. 20,000 parental or GD3S-KO RM-1 or 22Rv1 cells were plated in low serum medium in top chambers of trans-well chambers without (migration) or with (invasion) Matrigel coating and allowed to migrate or invade for 16 hours towards serum-containing medium in the bottom chambers. Left, representative images. Right, quantification. *n* = 3 with three replicates each and 6 images per well. (**I & J**) Reduced CRPC tumorsphere forming ability upon GD3S-KO. The indicated cell lines were plated in 24-well ultra-low attachment plates in 4% Matrigel and images were obtained on Day 7. Left, representative images; scale bar, 1000 µm. Right, quantified tumorsphere numbers per 4X microscopic field. n = 3 with three replicates each and 6 images per well. All Data represents mean ± SEM with unpaired *t* test; **, *P* < 0.01; ***, *P* < 0.001. ***p<0.001.

We also generated GD3S-KO derivative of the enzalutamide-resistant C4-2BER cell line, which has been reported to express NE markers [28] and expresses high GD2 levels compared to its parental cell line (**Supplementary Fig.S2C and S8A**). GD3S-KO eliminated the expression of GD2 in this model as well (**Supplementary Fig. S8B**). Furthermore, GD3S-KO led to reduction in the expression of NE differentiation markers, neuron specific enolase (NSE), Enhancer of Zeste homolog 2 (EZH2) and chromogranin A (CHGA) compared to C4-2BER cells, with levels in GD3S-KO cells comparable to those in GD2^low^ C4-2B cells (**Supplementary Fig.S8C**). These results suggest GD2 expression may regulate NE differentiation.

Functional analyses of various oncogenic traits showed a significant impact of GD3S-KO. Using CellTiterGlo assays, the GD3S-KO RM-1 cells exhibited a fold-increase of ∼51-fold compared to ∼73-fold for control cells, while GD3S-KO 22Rv1 cells showed a ∼18-fold-increase compared to a ∼28-fold increase for parental cells, with the differences being statistically significant (p <0.05) (**Fig. 4 E-F**). In cell migration assays, GD3S-KO RM1 and 22RV1 cell lines exhibited a 55% or 65% lower migration, respectively, relative to their controls (p<0.001) (**Fig. 4 G**). A similar impairment of trans-well invasion ability was seen upon GD3S-KO, with the GD3S-KO RM-1 and 22Rv1 cell showing 67.5 % and 73 % lower invasion, respectively, relative to their parental cell lines (p<0.001) (**Fig. 4 H**). Tumorsphere assays showed that GD3S-KO RM-1 and 22Rv1 cell lines exhibited 88% or 90% reduced tumorsphere forming abilities compared to their parental non-targeted cells (p<0.001) (**Fig. 5A**), supporting the importance of GD3S and its ganglioside products in promoting the CSC behavior of CRPCs. Further supporting this notion, FACS analysis for the proportion of CD49f^hi^ cells, a marker associated with PC CSCs [45], showed a reduction from 47% to 22% for GD3S-KO RM-1 (p<0.01) and 53% to 39% for GD3S-KO 22Rv1(p<0.05), relative to their parental lines (**Fig. 5 B**). To further evaluate the potential linkage of GD3S and GD2 expression with CSC characteristics, we analyzed the mRNA expression levels of 84 mouse genes included in a commercial CSC signature panel (RT^2^ profiler array) using quantitative PCR (qPCR) analysis of control vs. GD3S-KO RM-1 cells. The represented genes correspond to those associated with CSCs as well as Epithelial Mesenchymal Transition (EMT), given the strong mechanistic linkage of EMT and CSC behaviors [46]. Volcano plot analysis revealed multiple genes in the array to be either down-regulated or upregulated in the GD3S-KO derivative vs the control RM-1 cells (**Fig. 5C**). As presented in a heatmap (**Fig. 5D**), we observed a significant downregulation of 17 genes including *Notch1, Stat3*, *Sox2, Pou5f1(Oct4), Mycn, Lin28b, Snai1, Zeb1, Epcam, Pecam1* and upregulation of 22 genes such as *CD24a*, *Mertk, Slug*, *Foxa2*, *CD44*, *Aldha1a*, *Klf4* and *Myc* in GD3S-KO cells (**Fig. 5D**). Western blotting analysis validated the qPCR-observed downregulation of Snail, ZEB1, Lin28b, Notch1, Stat3 (⍺, β), SOX2 and the observed upregulation of Slug in GD3S-KO RM-1 and 22Rv1 cells vs. their control cells (**Fig. 5E**). Additional Western blot analysis of CSC/EMT markers SOX9 and N-cadherin revealed their downregulation in GD3S-KO RM-1 and 22Rv1 cells vs. their control cells (**Fig. 5E**). Collectively, these results are consistent with a conclusion that the GD2^high^ fraction in CRPC models defines a CSC-like population.

**Fig. 5:**
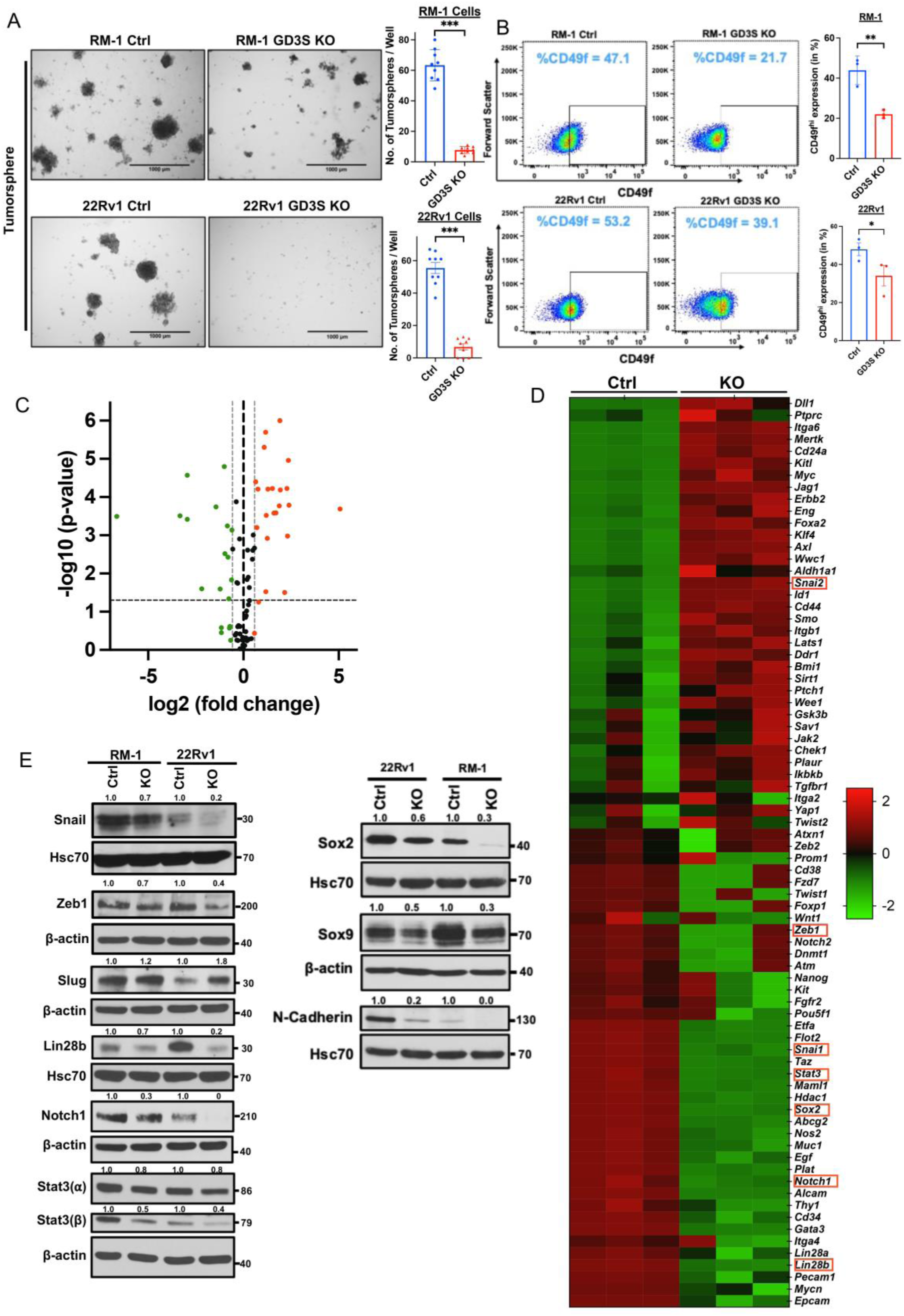
Impairment of cancer stem cell (CSC)-associated traits in CRPC cell lines upon GD3S-KO. (**A**) Reduced CRPC tumorsphere forming ability upon GD3S-KO. The indicated cell lines were plated in 24-well ultra-low attachment plates in 4% Matrigel and images were obtained on Day 7. Left, representative images; scale bar, 1000 µm. Right, quantified tumorsphere numbers per 4X microscopic field. *n* = 3 with three replicates each and 6 images per well **(B)** Reduced expression of CSCs markers in GD3S-KO CRPC cell lines. The indicated cell lines were subjected to live cell staining followed by Left, representative FACS plots display CD49f staining (MFI’s) on X-axis and forward scatter on Y-axis. Right, quantitation of % of CD49f-high population. *n*=3. (**C**) Volcano plot of the mouse cancer stem cell RT^2^ profiler quantitative PCR array analysis of RM-1 controls vs. GD3S-KO cells. X-axis, log fold change; Y-axis, -log p-value. Green dots represent the downregulated genes and red dots display the upregulated one with a log1.5 -fold change cut-off. (**D**) The heatmap shows the significantly upregulated (red) and downregulated (green) genes in RM-1 GD3S-KO cells compared to control cells.; *n*=3 independent experiments, p<0.05 is deemed significant. (**E**) Reduced expression of cancer stem cells and epithelial-mesenchymal transition (EMT) markers in GD3S-KO CRPC cell lines. Lysates of the parental vs. GD3S-KO RM-1 or 22Rv1 cell lines were subjected to western blotting with antibodies against the indicated proteins, with Hsc70 or β-actin used as loading controls. Densitometric quantification is shown on top of each blot. Data represents mean ± SEM, unpaired *t test*; *p<0.05, **p<0.01, ***p<0.001.

### GD3S-KO impairs the tumorigenic ability of GD2^high^ CRPC cell lines

To assess if the marked reduction in the *in vitro* pro-oncogenic traits of PC cell lines upon GD3S-KO translates into impaired tumorigenesis *in vivo,* we used lentiviral infection to introduce a tdTomato-luciferase dual reporter into control and GD3S-KO RM-1 cell lines (the latter with two distinct sgRNAs). The tdTomato-high fractions were enriched by FACS-sorting and stable cell lines selected. We implanted the reporter-bearing control and GD3S-KO RM-1 cells in the tibias of castrated syngeneic C57BL/6 mice and monitored tumor growth (5 mice/group) over time by bioluminescence imaging. While control RM-1 cell implants generated tumors that showed a time-dependent increase in log10 photon flux in IVIS imaging that peaked at 512-fold increase at the endpoint compared to day 7, both of the GD3S-KO RM-1 cell implanted groups generated significantly smaller tumors whose average peak photon flux was 40-fold (sgRNA1) and 20-fold (sgRNA2) lower compared to the control group (p<0.01 for both KO groups relative to control) (**Fig. 6A, B**). Morphometric analysis of tibial bone by micro-CT scanning showed reduced bone destruction by implants of GD3S-KO RM-1 cells relative to control as demonstrated by quantification of the bone volume (38.85% and 38.86% for sgRNA1 and sgRNA2 groups, respectively, relative to 30.5 % in control; p values for both <0.05), trabecular thickness (thickness of 141mm and 144 mm for sgRNA1 and sgRNA2 groups, respectively, relative to 127 mm in control; p values for both <0.05), trabecular number (trabecular number per mm of 0.00288 and 0.00278 for sgRNA1 and sgRNA2 groups, respectively, relative to 0.0023 in control; p values of <0.01 and <0.05 for sgRNA1 and sgRNA2 groups, respectively) and separation (516mm and 527mm for sgRNA1 and sgRNA2 groups, respectively, relative to 470 mm in control; p values for both <0.05) (**Fig. 6C-G**). To confirm the impact of GD3S-KO to impair the ability of implanted PC cells to form tumors in vivo, we further implanted the mCherry-luciferase expressing parental or GD3S-KO 22Rv1 cells intra-tibially in nude mice and analyzed these by IVIS imaging. Compared to a 15,984-fold increase in Log10 photon flux relative to time 0 for the control group, the GD3S-KO cell implants showed an increase of only 397.5-fold in peak Log10 photon flux (p<0.001) (**Fig. 6H, I**). Using IVIS, we did not observe distal metastases within the time of observation used in our studies (although some animals showed bioluminescent signals at the initial observation time point) (**Fig. 6H**). Together, these analyses demonstrate that GD3S-KO in CRPC cell line models significantly impairs their ability to form tumors when implanted in bone and to induce bone destruction.

**Fig. 6:**
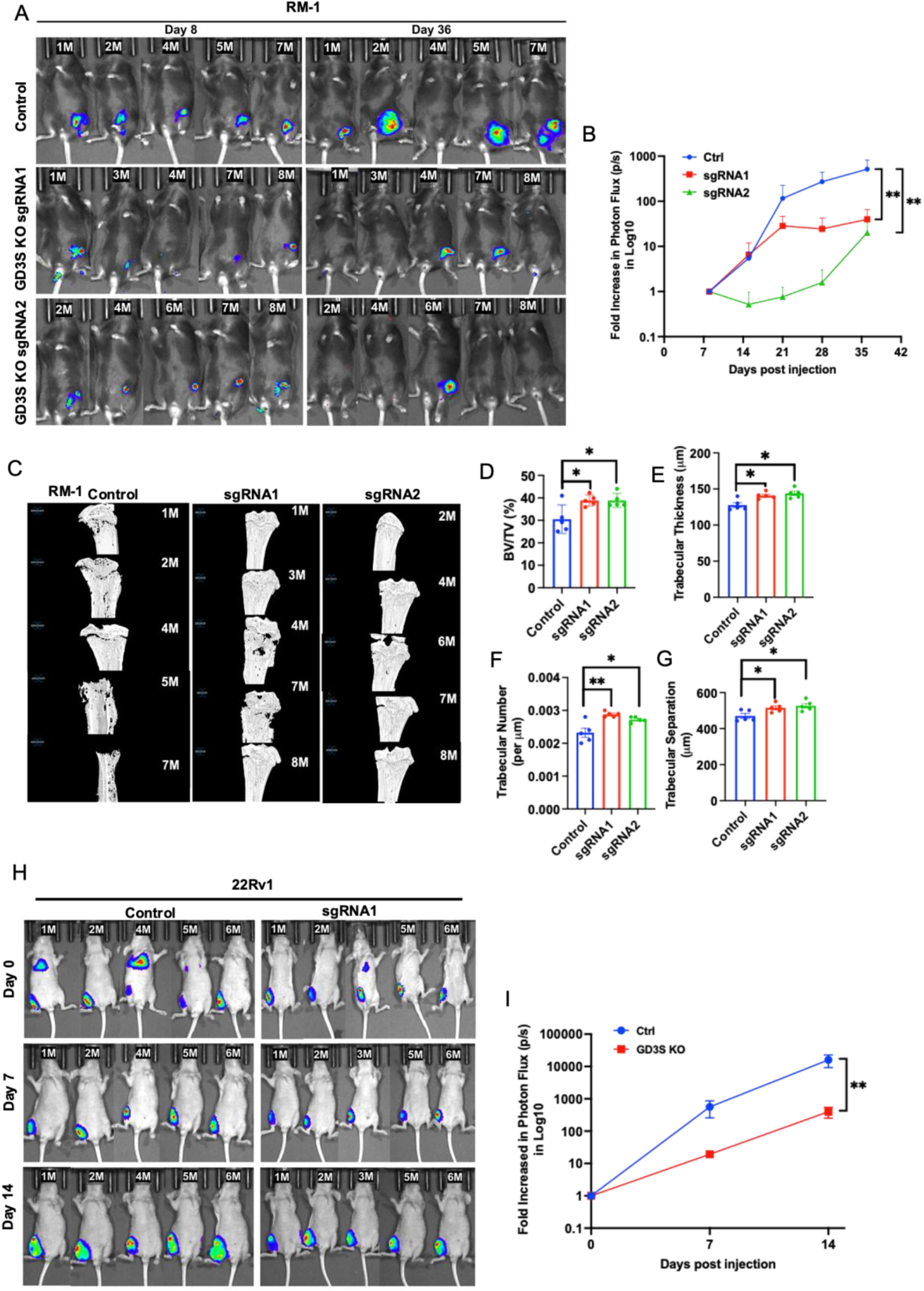
Impairment of *in vivo* bone-implanted CRPC tumorigenesis by GD3S-KO. (**A**) Parental (Control) RM-1 cells or their edited versions carrying sgRNA1 or sgRNA2 (GD3S-KO) were engineered with tdTomato-luciferase and 4 × 10^4^ cells injected in tibias of castrated male C57/BL6 mice (8/group) and primary tumor growth was monitored by bioluminescence imaging at the indicated time points in mice (5/group) with detectable bioluminescent signals on day 8. (**B**) Bioluminescence signals of RM-1 tumors over time are shown as log fold-change in photon flux over the time. Data are mean ± SEM, n=5; Two-way ANOVA (mixed model); *p<0.05. (**C**) Micro-CT scanned images of tibias isolated from mice implanted with the indicated RM-1 cell lines. μCT analyses of the trabecular bone was used to calculate the: (**D**) bone volume fraction (BV/TV%); (**E**) trabecular thickness (µm); (**F**) trabecular number (per µm); and (**G**) trabecular separation (µm). Data represent mean ± SEM, with unpaired t-test used for statistical analysis; *p<0.05 and ** p<0.01. (**H**) Impairment of tumorigenesis of 22Rv1 CRPC cells by GD3S-KO. 1 × 10^5^ control or GD3S-KO (sgRNA1) 22Rv1 cells engineered with mCherry-luciferase were injected in tibias of castrated male athymic nude mice (6/group) and tumor growth was monitored by bioluminescence imaging in mice (5/group) with detectable bioluminescent signals on day. (**I**) Bioluminescence signals of 22Rv1 xenografts over time are shown as log fold-change in photon flux over time. Data are mean ± SEM, n=5, with analysis by Two-way ANOVA (mixed model); **p<0.01.

## DISCUSSION

Despite improvements in diagnosis and management of early PC, progression to CRPC and metastatic disease represent lethal transitions responsible for high death burden from PC. Targetable molecular pathways that mark or contribute to these lethal states can open new therapeutic avenues in PC. Here, we use PC patient-derived tumor tissue analyses and cell line-based mechanistic approaches to demonstrate that the di-ganglioside GD2 marks a small but functionally-critical tumor cell subpopulation in PC with enrichment in more advanced disease states. Our studies using CRPC cell models establish that GD2 defines a cancer stem cell-like subpopulation that contributes to oncogenic drive, thus raising the prospect of GD2 targeting in CRPC using clinically approved therapeutic agents.

While expression and functional roles of GD2 are established in a number of cancers, with GD2 targeting with antibodies now approved for therapy of neuroblastoma patients [8], only few studies have examined the expression of GD2 in PC and its roles are not known. No prior studies have examined GD2 expression in PC tissues at a histological level but limited prior studies primarily using biochemical fractionation of glycolipids [22-24] are consistent with our findings. The latter studies utilized a small number of normal or tumor tissues with biochemical analyses of GD2 expression which did not provide an assessment of what proportion of PC patients express GD2, and whether GD2 was uniformly expressed or in a heterogenous pattern. Our analyses using immunohistochemical staining (**Fig. 2**) therefore provide unique new insights by demonstrating that GD2 is expressed on a larger proportion of tumors versus the normal prostatic tissue, and importantly that only a small subset of cells in patient tumors express GD2. Notably, a significantly higher proportion of metastatic tumors harbored such cells (**Fig. 2E**). The GD2 expression pattern in PC is reminiscent of non-neuroectodermal tumors like breast cancer with a small GD2^+^ tumor cell subpopulation [9], with GD2 marking the rare stem cell population in normal tissues and tumors [47].

Our studies using cell line models and FACS analyses (**Table 1, Supplementary Fig. S2**) establish that similar to other well-studied tumor models, that GD2 is expressed at the cell surface of tumor cells. While even GD2^+^ cell line models showed a non-homogenous GD2 expression, similar to breast and other non-neuroectodermal tumor systems [9], the GD2^high^ and GD2^low^ populations appear to be in a dynamic equilibrium, as seen by the re-emergence of mixed GD2^high^/GD2^low^ populations when sorted GD2^high^ or GD2^low^ subsets were cultured *in vitro* (**Supplementary Fig. S4**), similar to that described in other tumor systems [41]. Whether GD2 expression in vivo is similarly dynamic is currently unknown, but potentially significant to explore in the future. In this regard, dramatic induction/elevation of GD2 expression under scenarios of significance to the evolution of CRPC, including experimentally promoting lineage plasticity of AR-driven C4-2 cells by TP53 or RB1-KD (**Fig. 1C-D**), or when GD2^low^ C4-2B cells were made enzalutamide-resistant (**Table 1, Supplementary Fig. S2**), suggest that GD2 may be functionally important in more advanced stages of PC, a suggestion consistent with our finding of high GD3S (*St8SIA1*) mRNA levels a subset of CRPC PDX and tumor-derived organoid models (**Supplementary Fig. S3)** represented in publicly-available transcriptomic data [26].

Notably, the IHC-based GD2 staining in PC tissues (**Fig. 2A-C**) as well as cell lines (**Fig. 1D & Supplementary Fig. S1C**) was predominantly intracellular (cytoplasmic) in contrast with the plasma membrane staining of cell lines analyzed by FACS (**Fig. 3D & E; Supplementary Fig S2**). Published IHC studies showed a similar patten of GD2, described as perinuclear granular “Golgi-like” pattern, in neuroblastoma [48] and breast cancer tissues [49]. It is likely that the difference reflects the use of live cells for FACS analysis (precluding any staining of intracellular pools of GD2) while IHC is performed on fixed and permeabilized tissue sections where intracellular GD2 would be accessible to the staining antibody.

That GD2 expression on only a subset of tumor cells was nonetheless critical for tumorigenesis (**Fig. 6**) is reminiscent of studies in breast and other cancers where depletion of GD2 by genetic or pharmacologic downregulation of GD3S expression [11, 50] or by immunologic targeting of cell surface GD2 [12] led to impaired tumorigenesis. In these previous studies, GD2 expression marked the CSC population, thus accounting for GD2’s role in multiple tumorigenic traits. Strong co-staining of GD2 with the CSC marker, high CD49f, enrichment of the GD2^+^ (**Fig. 3G**) and CD49f^hi^ (CSC) populations (**Fig. 5B**) in tumorspheres, the requirements of GD3S for tumorsphere growth (**Fig. 5A**) and alterations in CSC and EMT-related signature gene expression upon GD3S-KO (**Fig. 5C-E**) are consistent with this idea. However, more in-depth analyses of the GD2^high^ and GD2^low^ subpopulations of PC models, including PDX and organoid models expressing GD2, examining their tumor-initiation and maintenance capabilities under limiting dilution conditions [9], resistance to conventional therapies (such as chemotherapy) [20] and other CSC behaviors will be needed to firmly establish that GD2^+^ subpopulation in PCs represents CSCs.

Since our current assignment of a functional role to GD2 relied on GD3S-KO, similar to GD3S depletion approaches employed by others [11, 50], additional studies are needed to formally rule out a role of GD3, the other GD3S product and precursor to GD2 [2]. Although the surface GD3 levels on the two cell models we targeted for GD3S-KO were relatively modest (**Supplementary Fig. S6 and S7**), further immunological targeting of GD2 and/or GD2S deletion should elucidate the functional role of GD2 vs. GD3 in PC tumorigenesis.

In our studies, we primarily utilized the antibody staining to demonstrate the expression of GD2 in cell lines and patient tumors. While this approach is well-established, and we further authenticated the antibody used (14G2a) in our studies (**Supplementary Fig. S1**), it is known that this antibody may also recognize modified forms of GD2, such as O-Acetyl-GD2 [51], that are expressed in some tumors. In limited analyses, using TLC separation of extracted lipids and standard GD2, we found the 14G2a to recognize a sphingolipid species comigrating with authentic GD2, and loss of this species upon GD3S-KO (**Supplementary Fig. S5**). However, further biochemical validation using TLC and/or Mass Spectrometry methods will be needed to establish that GD2 expression we define here in PC represents GD2 and not its modified forms. This is particularly important for future analyses of tumor tissues to avoid possible false positive/negative results as the tissue fixation and other processing could affect how the GD2 antigen is displayed in the samples.

Collectively, our novel findings reveal that GD2 overexpression is a feature of a large subset of PCs, that GD2 may be further induced during transitions associated with PC progression, and that the GD2 biosynthetic machinery enzyme GD3S is essential for efficient tumorigenesis in CRPC models. Our findings raise the prospect of anti-GD2 targeting in advanced PC.

## ACKNOWLEDGEMENTS

We thank Dr. Ian Frew for Multiple Lentiviral Expression System Kit and Dr. Irmela Jeremias for the pCDH-EF1a-eFFLy-mCherry (mCherry-luciferase reporter) plasmid, obtained through Addgene.

---

**Fig. S1.**
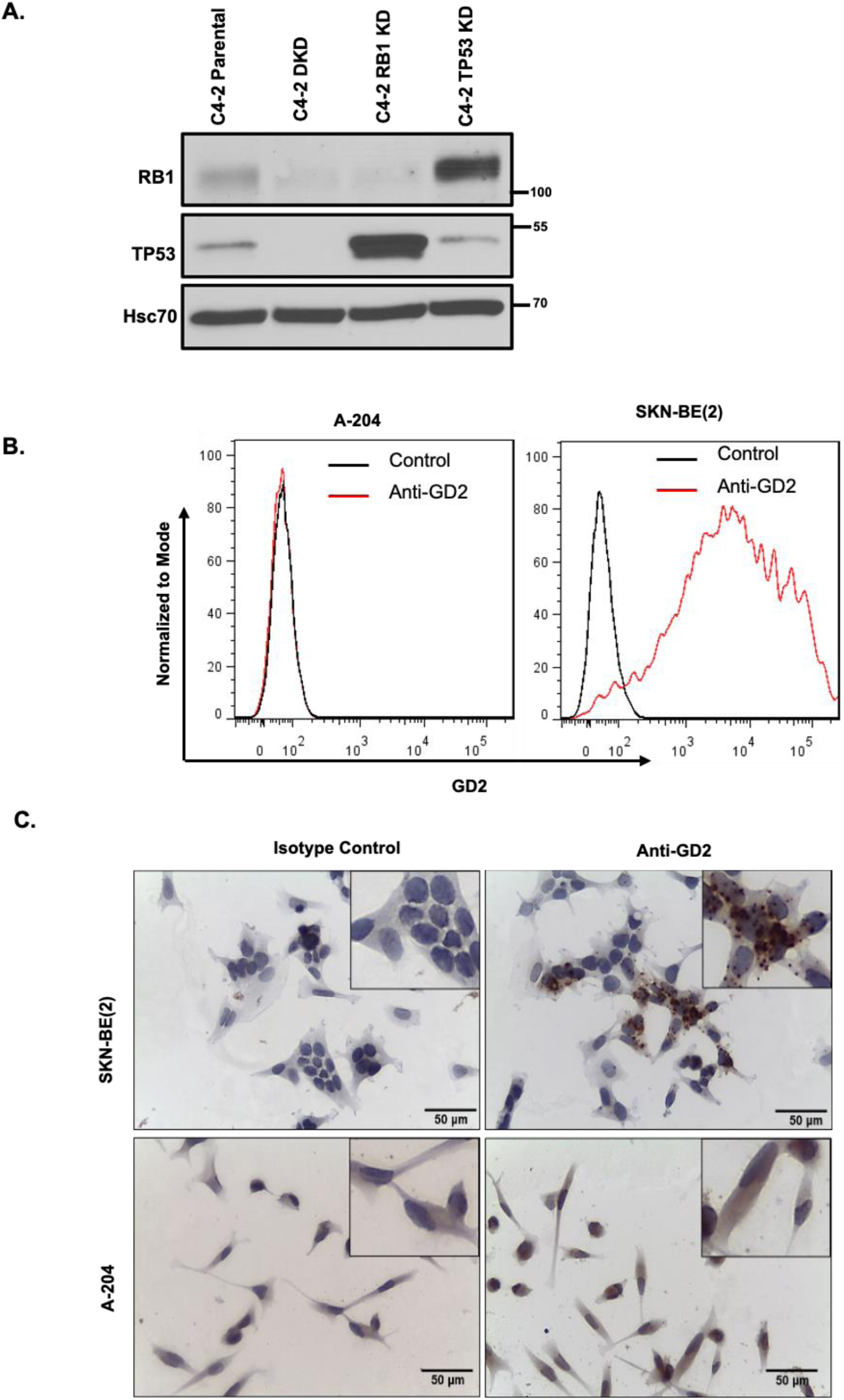
Confirmation of RB1, TP53 KD and RB1/TP53 double KD (DKD) in individual and DKD variants of C4-2 and validation of anti-GD2 antibody. **(A)** Western blotting shows the reduction of RB1, TP53 in respective knockdown cells and both RB1 and TP53 KD in DKD samples using indicated antibodies. HSC-70 used as a loading control. **(B)** FACS analysis of GD2^-^ A-204 Rhabdomyosarcoma and GD2^+^ SK-N-BE(2) human neuroblastoma cell lines stained with anti-GD2 vs. isotype control. **(C)** Immunohistochemistry (IHC) staining of A-204 and SK-N-BE(2) cell lines with anti-GD2 or isotype control. Scale bar, 50 µm. Insets show a higher magnification of regions within each image. GD2 antibody selectively stains the SK-NB-2 but not the A-204 cell line.

**Fig. S2:**
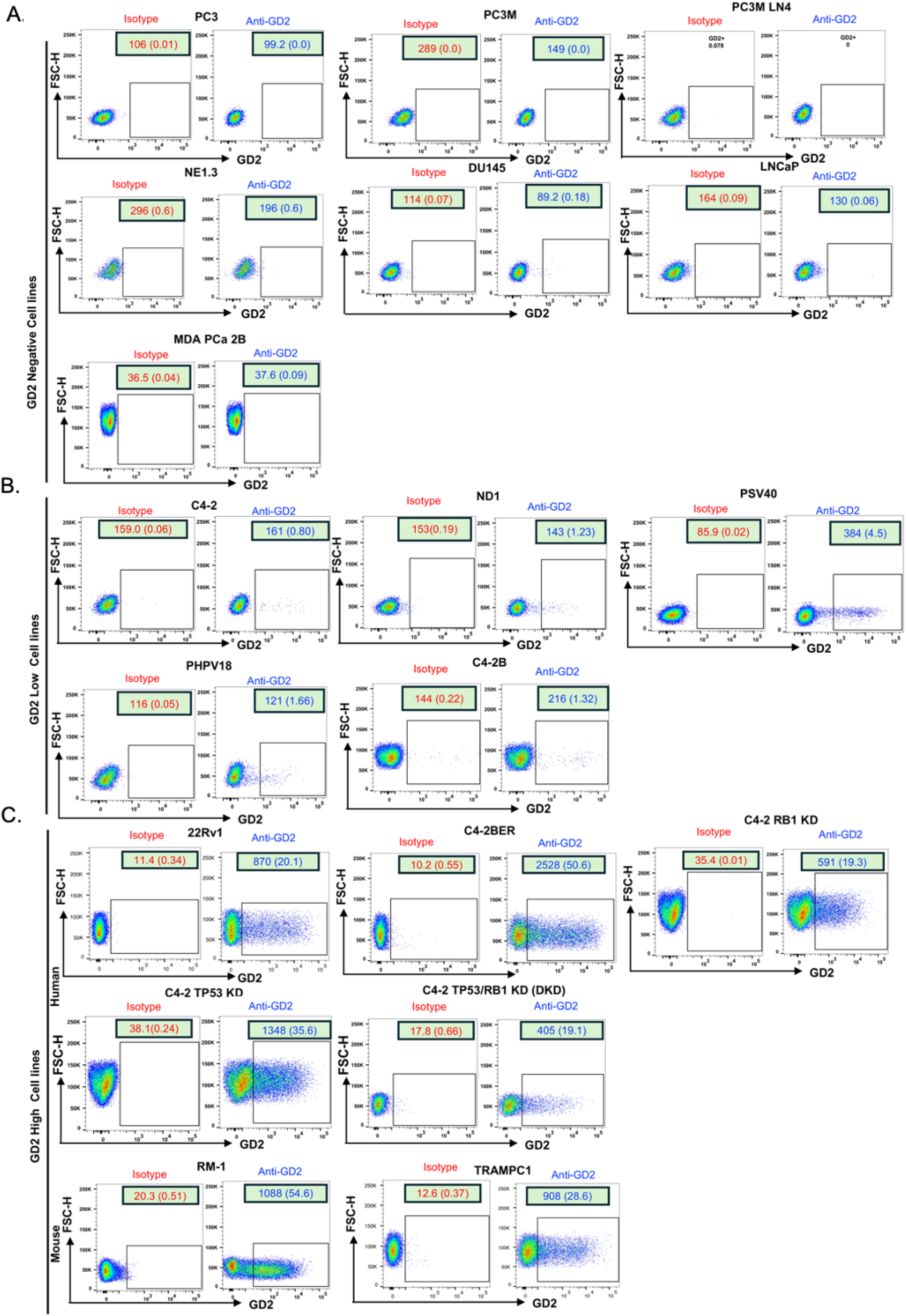
FACS plots of cell surface GD2 expression on human and mouse prostate cancer (PC) cell lines. The indicated human and mouse prostate cancer cell lines were live cell stained using anti-GD2 vs. isotype control and analyzed by FACS. GD2, blue; isotype control, red. Percentage % population and MFI are indicated in FACS dot plots. (**A**) Cell lines with essentially undetectable GD2 (all human). (**B**) GD2^low^ PC cell lines (all human). (**C**) GD2^high^ human (22Rv1 and C4-2BER) and mouse PC cell lines (RM-1 and Tramp-C1) are shown.

**Fig. S3:**
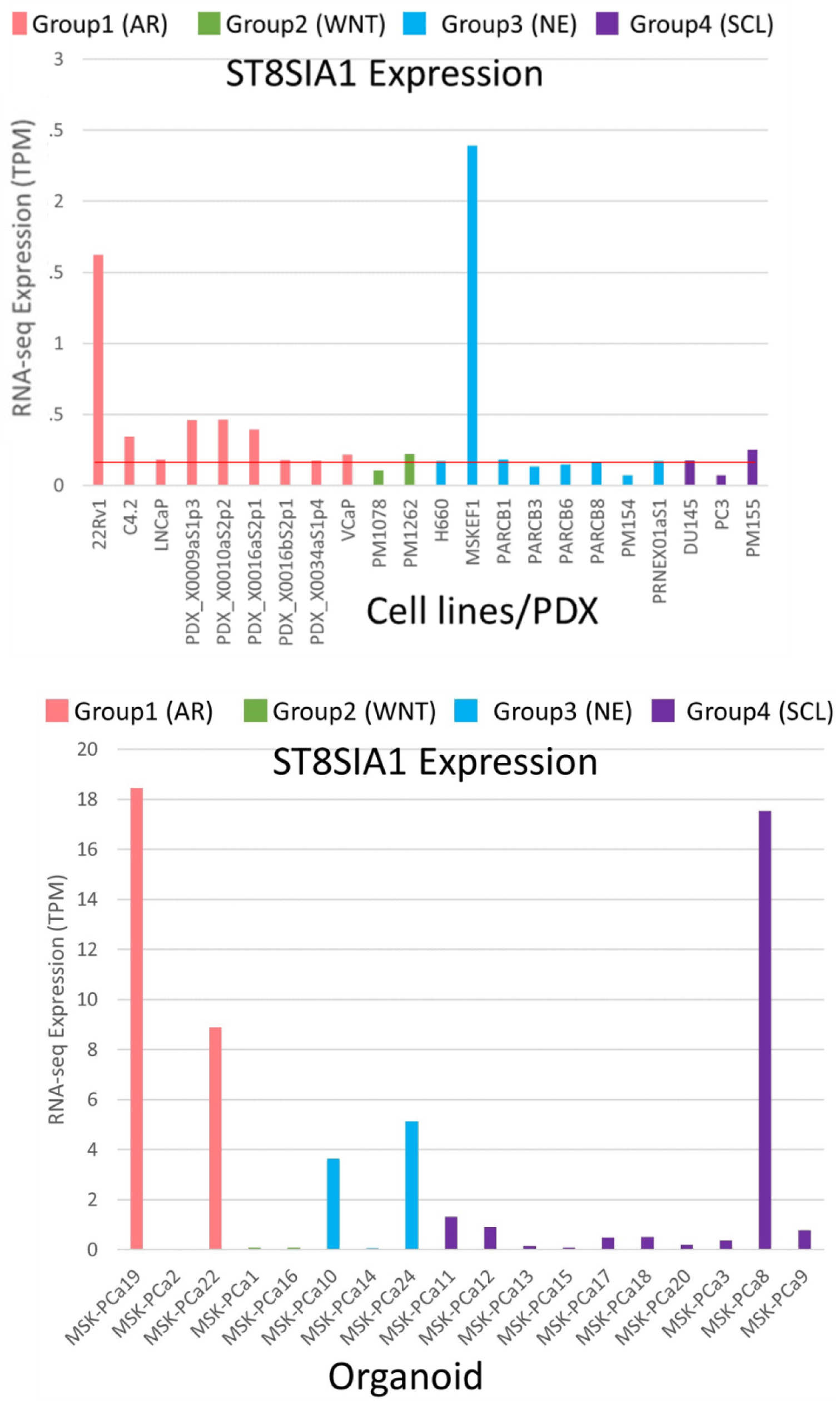
High *ST8SIA1* (GD3S) mRNA expression in a substantial proportion of CRPC patient-derived xenograft (PDX) tumors and organoid models. Publicly available transcriptomic data published by Tang et al.[1] (GEO accession GSE199190) on CRPC PDX and cell lines (Top panel) and organoid models (Bottom panel) were queried for *ST8SIA1* mRNA expression. The CRPC groups are as per Tang et al: Group 1, Androgen Receptor dependent (CRPC-AR, peach); Group 2, Wnt dependent (CRPC-WNT, green); Group 3, neuroendocrine (CRPC-NE, blue); and Group 4, stem cell like (CRPC-SCL, purple). Gene expression is shown as normalized transcripts per million (TPM). The horizontal red line in left panel demarcates the negative/positive cutoff based on GD2^-^ LNCaP cells.

**Fig. S4:**
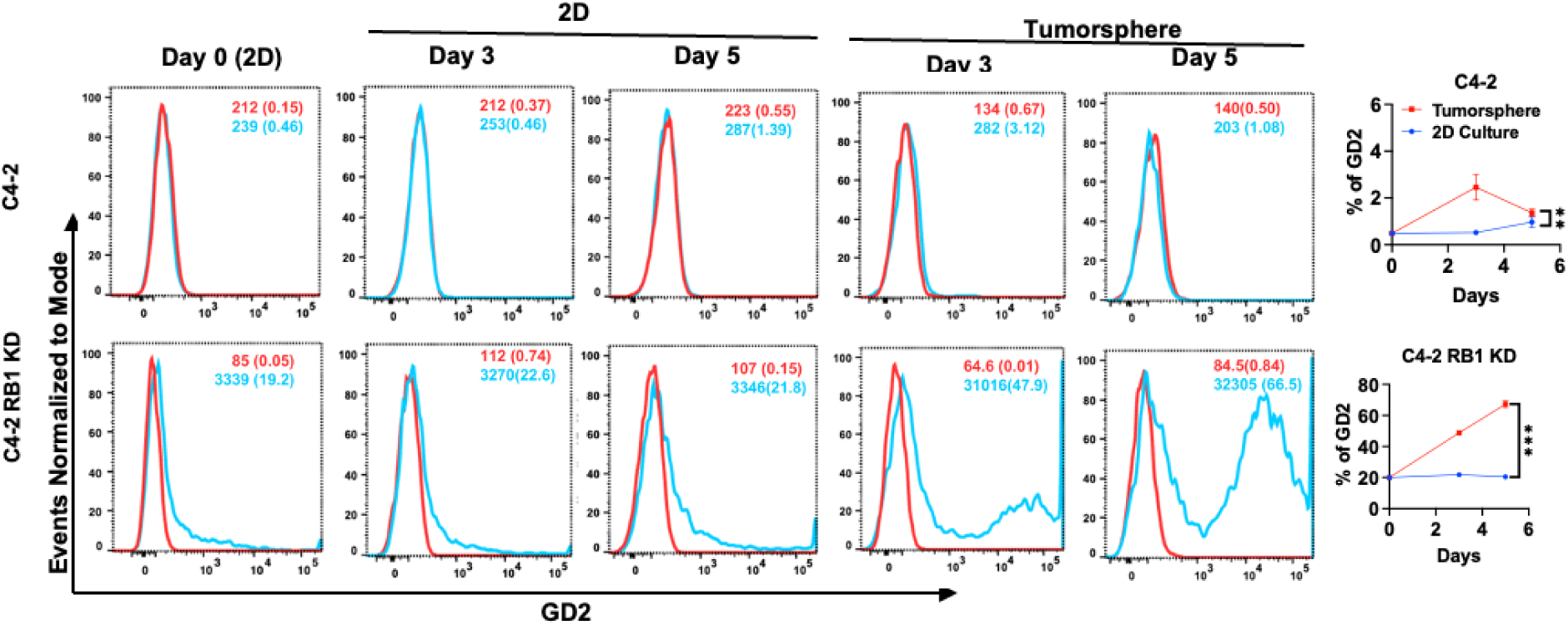
Enrichment of GD2^high^ cells in tumorspheres of CRPC cell lines: Enrichment of GD2^high^ cells in tumorspheres of CRPC cell lines. The C4-2 parental and RB1 KD cells were cultured in regular two-dimensional (2D) cultures or were seeded in tumorsphere cultures. Cells harvested at the indicated times were analyzed for cell surface GD2 expression (vs. isotype control) at the indicated times. % GD2^+^ cells and MFI are indicated. **Left**, representative FACS images; **Right**, quantitation of data. Mean ± SEM with two-way ANOVA test, **, *P* < 0.01; ***, *P* < 0.001.

**Fig. S5:**
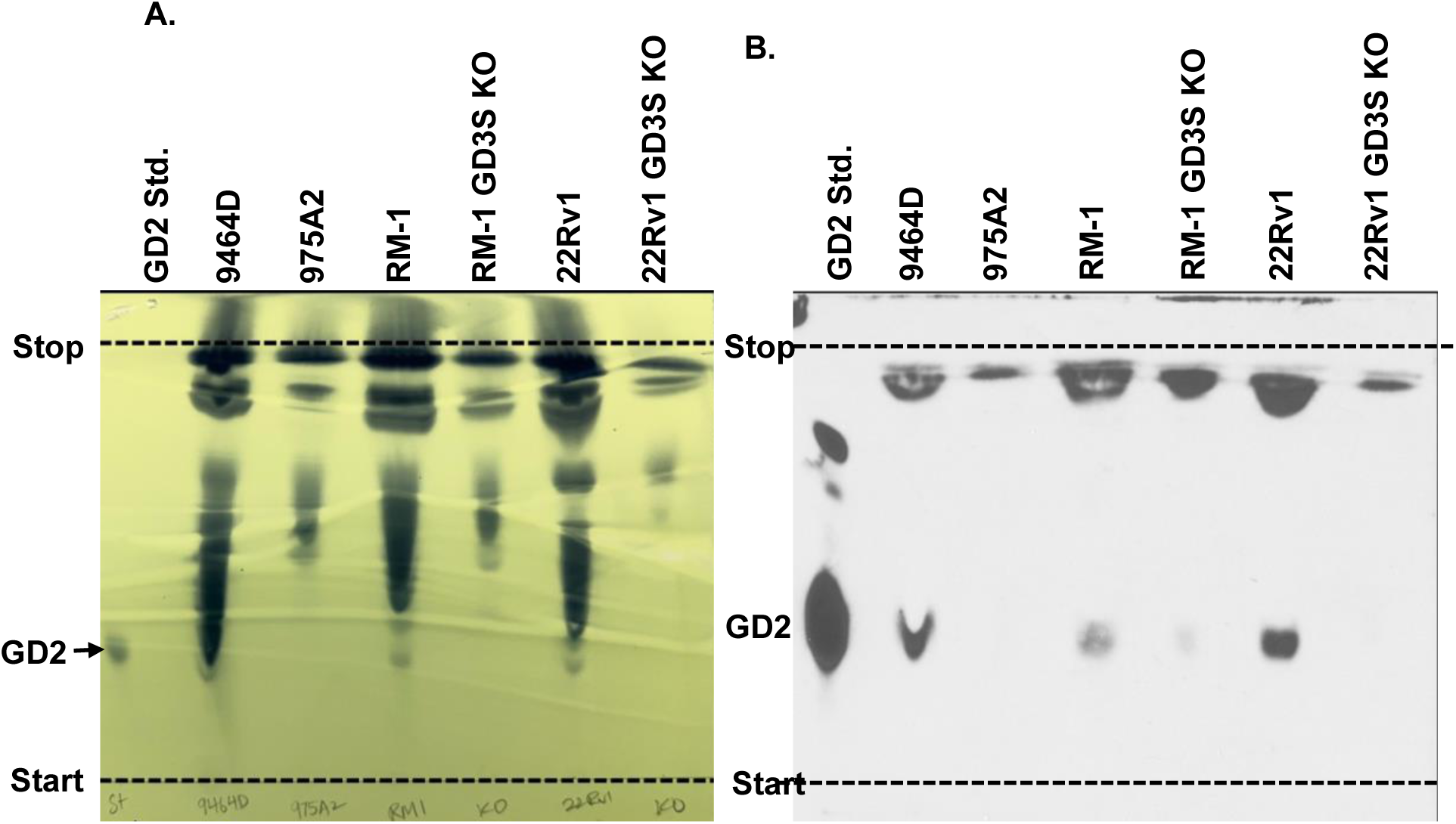
Thin Layer Chromatography (TLC) immunoblotting using ganglioside extracts demonstrate presence of GD2 in parental RM-1 and 22Rv1 cells and loss in GD3S KO cells. **(A)** Isolated gangliosides from indicated cell lines were resolved for TLC in silica plates. Mouse neuroblastoma cell lines, 9464D and 975A2 were used as positive and negative control for GD2 respectively. Purified GD2 obtained commercially was used as a positive standard. Represented Phosphomolybdic acid (PMA) staining of the resolved TLC plate show migration of GD2 in standard and faint comigration of GD2 in RM1, 22Rv1 and positive control 9464D cells and absence in GD3S KO and GD2 negative lanes. **(B)** TLC plate run in parallel with indicated cell lines were processed for immunoblotting with anti-GD2 antibody. GD2 immunoreactivity was observed in GD2 standard along with parental RM-1, 22Rv1 and positive control cell line 9464D.

**Fig. S6:**
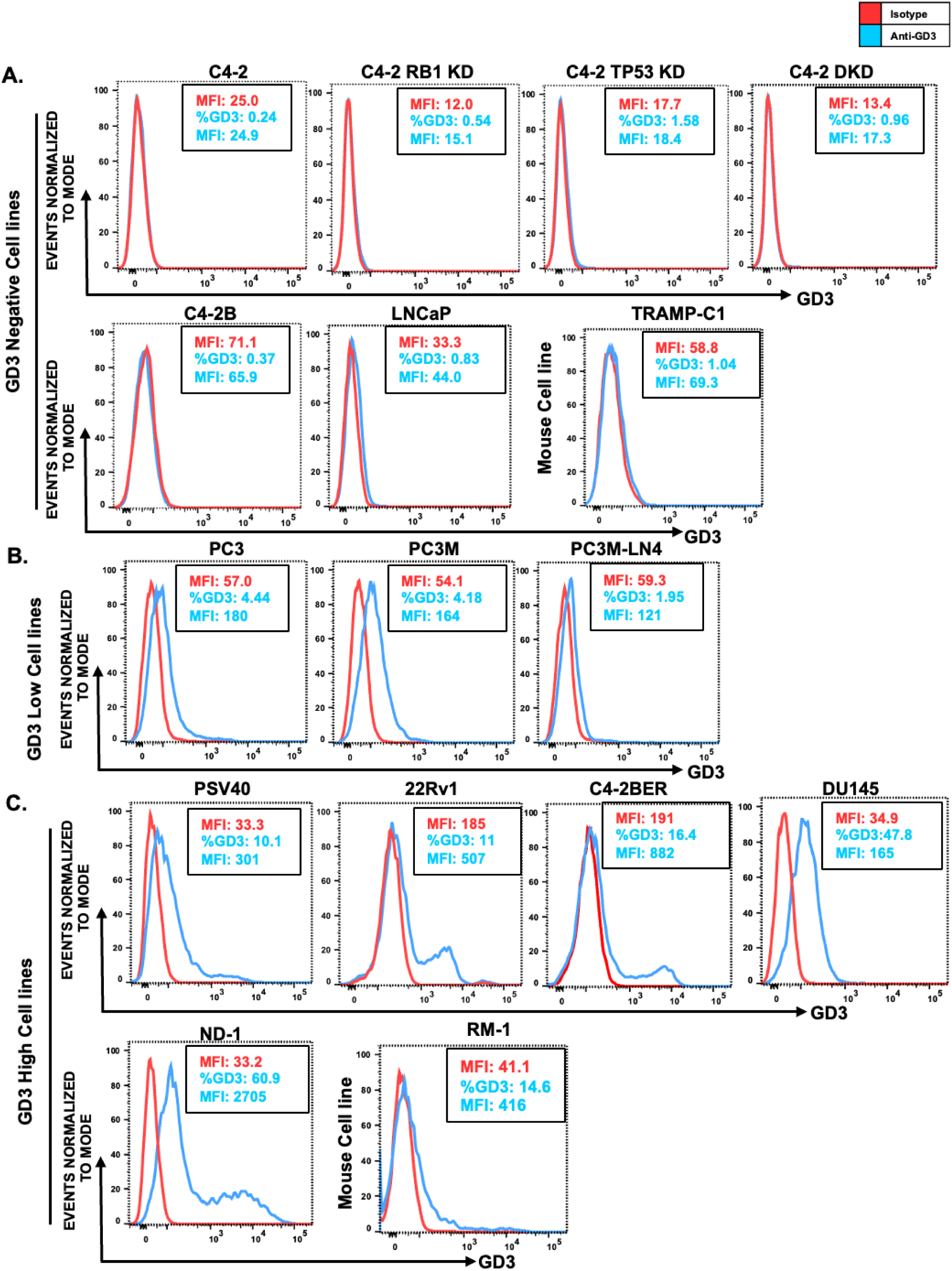
FACS plots of cell surface GD3 expression on human and mouse prostate cancer (PC) cell lines. The indicated human and mouse prostate cancer cell lines were live cell stained using anti-GD3 vs. isotype control and analyzed by FACS. GD3, blue; isotype control, red. Percentage % population and MFI are indicated in FACS dot plots. (**A**) Cell lines with essentially undetectable GD3. (**B**) GD3^low^ PC cell lines (all human). (**C**) GD3^high^ human (PSV40, 22Rv1, C4-2BER, DU145, ND-1) and mouse PC cell lines (RM-) are shown.

**Fig. S7:**
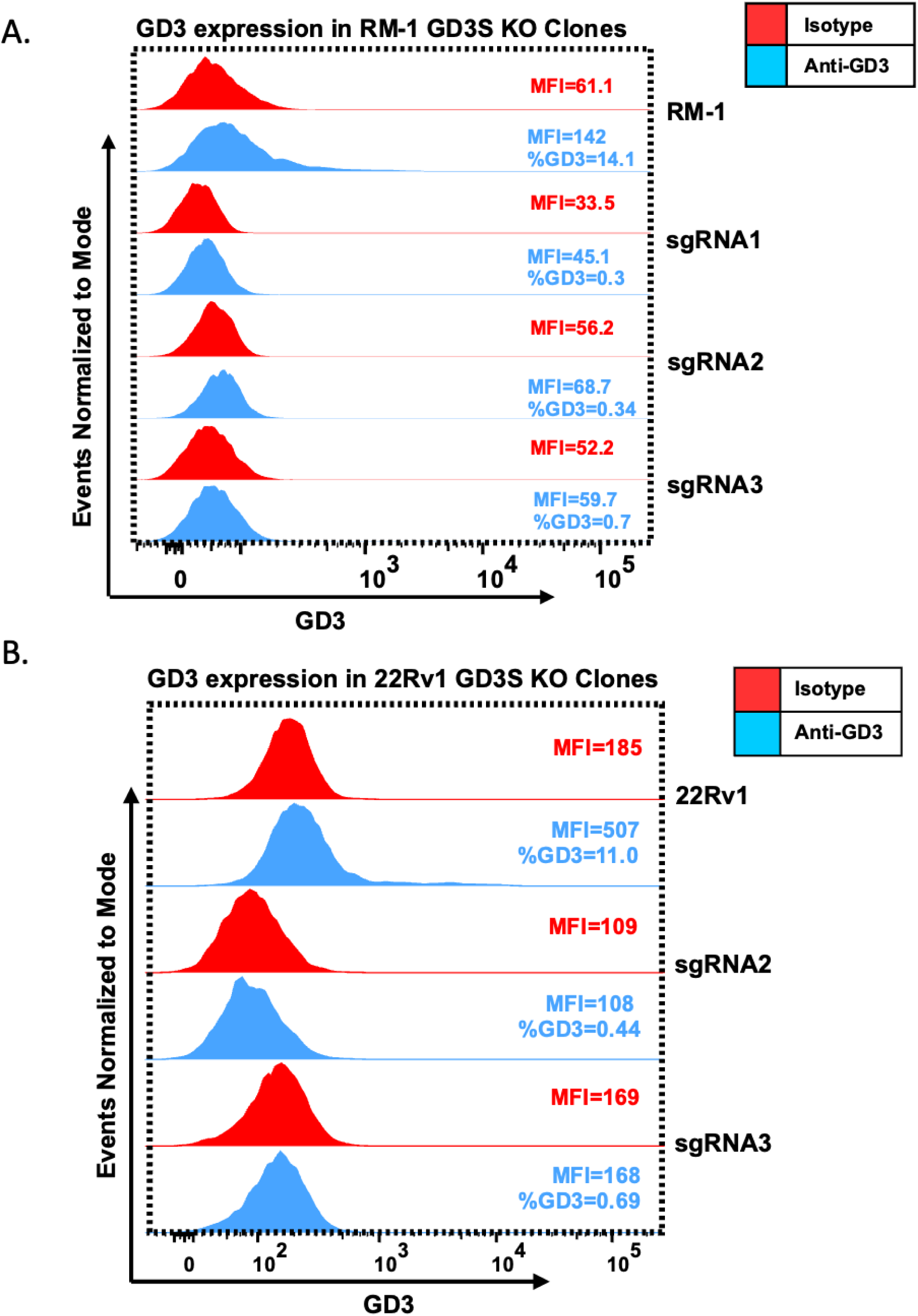
GD3 expression in GD3S knockout RM-1 cells. Live cell FACS analysis of murine RM-1 and its GD3S KO variants (sgRNA1, sgRNA2 & sgRNA3) **(A)** and human 22Rv1 and its GD3S KO variants (sgRNA2 & sgRNA3) **(B)** show decreased expression of GD3 synthesis in the GD3S KO variants and is displayed as over lay FACS plots. The percentage of GD3 and MFI are indicated in FACS plots. The names of the cell lines and antibodies are mentioned on the right side of the FACS plots.

**Fig. S8:**
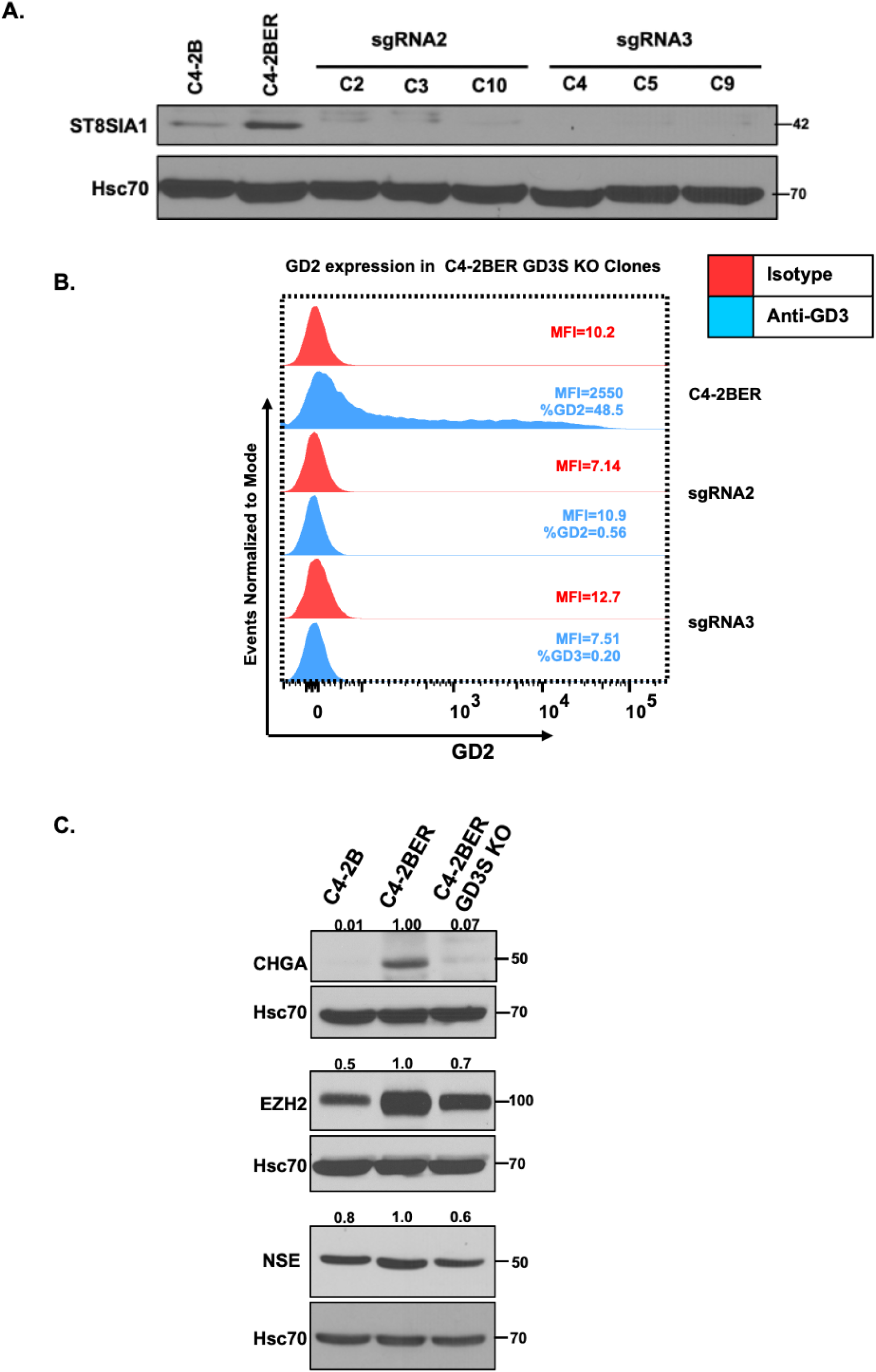
GD3S KO in C4-2BER cells demonstrate reduction of neuroendocrine markers: **(A)** Generation of GD3S knockout of RM-1 and 22Rv1 CRPC cell lines. Cells were transduced with lentiviral All-in-One CRISPR/Cas9 constructs and stable clones analyzed by Western blotting with anti-GD3S antibody; Hsc70, loading control. **(B)** Live cell FACS analysis of human C4-2BER and its GD3S KO variants (sgRNA2 & sgRNA3) show decreased expression of GD2 in the GD3S KO variants and is displayed as over lay FACS plots. The percentage of GD2 and MFI are indicated in FACS plots. The names of the cell lines and antibodies are mentioned on the right side of the FACS plots. **(C)** Representative Western blotting shows the increased neuroendocrine (NE) differentiation markers CHGA, EZH2 and NSE in enzalutamide resistant C4-2BER cells and loss of upregulated NE markers in GD3S KO variant of C4-2BER. Densitometries are indicated on the top of the bands after normalizing with HSC-70. Three independent experiments were performed.

